# Identification of a plant kinase that phosphorylates the bacterial effector AvrPtoB

**DOI:** 10.1101/2020.03.21.001826

**Authors:** Lei Lei, Danielle M. Stevens, Gitta Coaker

**Affiliations:** Department of Plant Pathology, University of California, Davis, Davis, CA, USA

## Abstract

A critical component controlling bacterial virulence is the delivery of pathogen effectors into plant cells during infection. Effectors alter host metabolism and immunity for pathogen benefit. Multiple effectors are phosphorylated by host kinases, and this posttranslational modification is important for their activity. We sought to identify host kinases involved in effector phosphorylation. Multiple phosphorylated effector residues matched the proposed consensus motif for the plant calcium-dependent protein kinase (CDPK) and Snf1-related kinase (SnRK) superfamily. The conserved *Pseudomonas* effector AvrPtoB acts as an E3 ubiquitin ligase and promotes bacterial virulence. We identified a member of the Arabidopsis SnRK family, SnRK2.8, which associated with AvrPtoB in yeast and *in planta*. *SnRK2.8* was required for AvrPtoB virulence functions, including facilitating bacterial colonization, suppression of callose deposition, and targeting the plant defense regulator NPR1 and flagellin receptor FLS2. Mass spectrometry revealed AvrPtoB phosphorylation at multiple serine residues *in planta*, with S258 phosphorylation significantly reduced in the *snrk2.8* knockout. AvrPtoB phospho-null mutants exhibited compromised virulence functions and were unable to suppress NPR1 accumulation, FLS2 accumulation, or inhibit FLS2-BAK1 complex formation upon flagellin perception. These data identify a conserved plant kinase utilized by a pathogen effector to promote disease.

## Introduction

Plants are exposed to diverse pathogens and rely on both passive and active defenses in order to restrict infection. Passive plant defenses include a waxy cuticle, pre-formed antimicrobial compounds, and the cell wall (Gu et al., 2017). Inducible defenses are often triggered by membrane-localized pattern recognition receptors (PRRs) as well as intracellular nucleotide-binding site leucine-rich repeat proteins (NLRs) (Boutrot and Zipfel, 2017; Lolle et al., 2020). Common plant immune responses include the production of reactive oxygen species (ROS), callose deposition, activation of mitogen-activated protein kinases, and global transcriptional reprogramming towards defense (Bigeard et al., 2015; Peng et al., 2018).

Pathogens secrete effector proteins that act to suppress plant immune responses, alter host developmental processes, and affect host metabolism to promote pathogen infection (Toruño et al., 2015). Despite the collective importance of effectors for pathogen virulence, higher-order effector deletions are often required to observe strong effects. However, two *P. syringae* effectors, *AvrPto* and *AvrPtoB*, substantially increase bacterial virulence (Cunnac et al., 2011). AvrPto is a small effector that acts as a kinase inhibitor, while AvrPtoB is a large, multidomain protein. Both effectors target similar plant proteins and therefore are functionally redundant with respect to bacterial virulence (Abramovitch et al., 2006; Xiang et al., 2008). Both AvrPto and AvrPtoB associate with the flagellin receptor FLAGELLIN SENSING 2 (FLS2) and its co-receptor BRASSINOSTEROID INSENSITIVE 1-ASSOCIATED RECEPTOR KINASE 1 (BAK1) and disrupt flagellin 22 (flg22) induced FLS2-BAK1 complex formation (Göhre et al., 2008; Shan et al., 2008; Xiang et al., 2008). The N-terminus of AvrPtoB comprises multiple target-binding domains and its C-terminus is a U-box type E3 ubiquitin ligase (Xiao et al., 2007a; Abramovitch et al., 2006). AvrPtoB mediates the degradation of multiple plant components, including FLS2, the salicylic acid (SA) defense pathway regulator NON-EXPRESSER OF PR GENES 1 (NPR1), the kinase Fen, the chitin receptor CERK1, and the exocyst complex protein EXO70B1 (Rosebrock et al., 2007; Gimenez-Ibanez et al., 2009; Chen at al.,2017; Wang et al., 2019). AvrPtoB’s diverse targets demonstrate its prominent role in *P. syringae* infection.

Effectors are produced in the pathogen but function inside their host. Several pathogen effectors rely upon host phosphorylation to promote their virulence activity (Xing et al., 2002; Bhattacharjee et al., 2015). Truncated AvrPtoB1-307 is phosphorylated in different plants including tomato, *Nicotiana benthamiana*, and *Arabidopsis thaliana*. The substitution of the serine 258 phosphorylated residue on AvrPtoB_1-307_ to alanine results in significantly attenuated virulence on susceptible tomato genotypes (Xiao et al., 2007b). The *P. syringae* effectors AvrPto, AvrB, and HopQ1 as well as the *Rhizobium* effectors NopL, NopP, and the cyst nematode effector 10A07 also target and recruit host kinases to promote their virulence (Skorpil et al., 2005; Anderson et al., 2006; Desveaux et al., 2007; Yeam et al., 2009; Zhang et al., 2011; Li et al., 2013; Hewezi et al., 2015). However, except for the nematode effector 10A07, host kinases utilized by these pathogen effectors remain unknown. In this study, we investigated the identity of the host kinase that phosphorylates AvrPtoB. Our results identify a conserved plant kinase recruited by this core *P. syringae* effector to promote virulence.

## Results and Discussion

### The *Pseudomonas syringae* effector AvrPtoB interacts with the plant kinase SnRK2.8

To investigate which plant kinases may be capable of phosphorylating pathogen effectors, we analyzed the sequence of previously identified effector phosphorylation sites (Supplemental Table 1). Interestingly, most effector phosphorylation sites, including AvrPtoB’s phosphorylated residues, matched the proposed consensus phosphorylation motif of the sucrose non-fermenting-1 (SNF1)-related kinases (SnRKs) and calcium-dependent protein kinases (CDPKs) (R-X_(2-3)_-S/T or S/T-X_(1-2)_-R) (Klimecka and Muszyńska, 2007; Vlad et al., 2008) (Supplemental Figure 1). The SnRK-CDPK family is conserved in plants and involved in a range of metabolic and stress signaling pathways, such as carbohydrate biosynthesis, abscisic acid (ABA)-induced signaling, salinity tolerance, cold stress, and response to pathogen infection (Coello et al., 2011; Hulsmans et al., 2016). The plant SnRK family can be subdivided into three subfamilies: SnRK1, SnRK2, and SnRK3. Multiple SnRK members are transcriptionally induced upon activation of plant defense, including application of SA, the immunogenic flagellin epitope flg22, or infection with the *P. syringae* type III secretion mutant ∆hrcC (Figure 1A).

**Figure 1.**
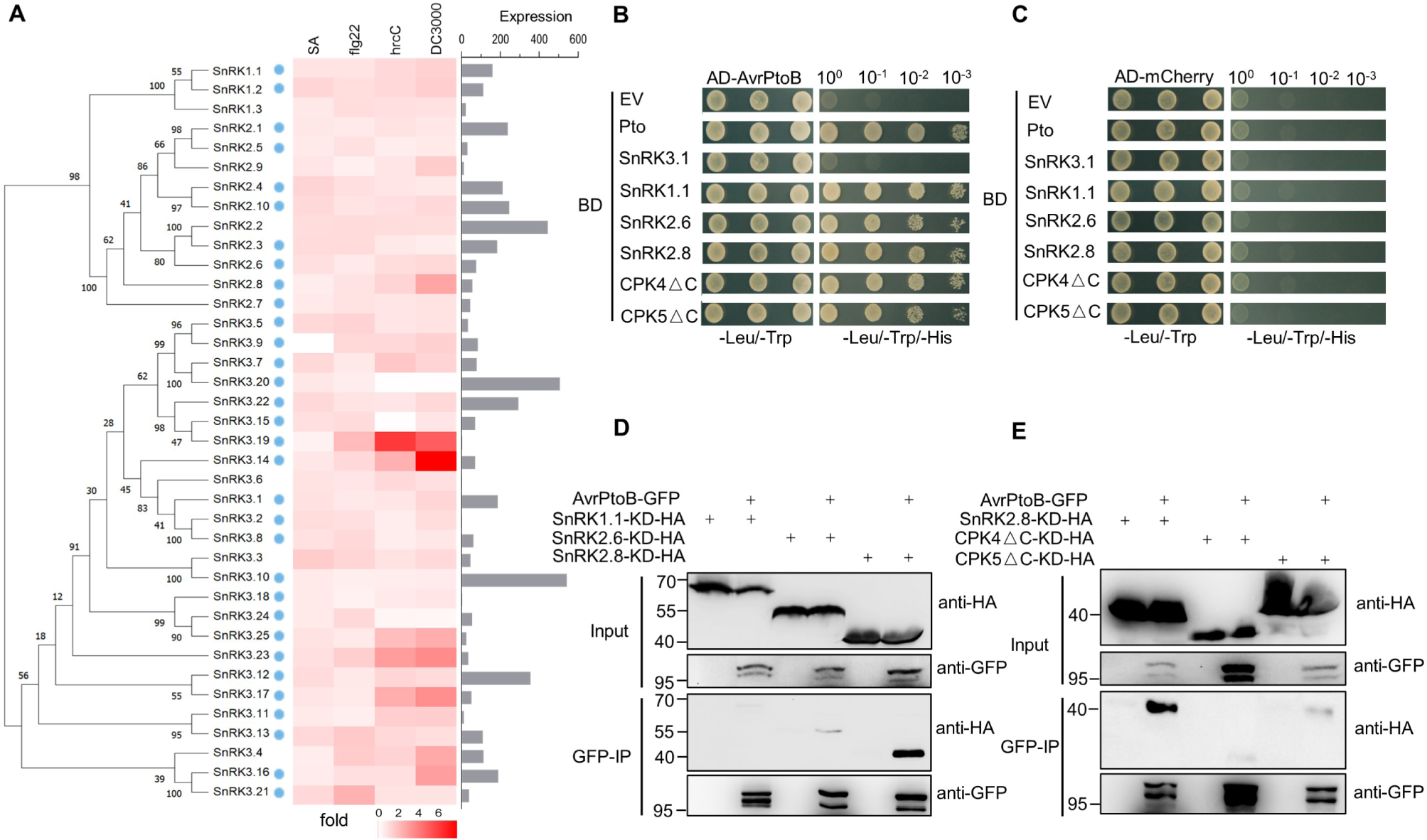
The *Pseudomonas syringae* effector AvrPtoB interacts with the plant kinase SnRK2.8. (A) Phylogeny of the *Arabidopsis* SnRK family and transcript expression of *SnRK* members after immune activation and pathogen infection. Phylogeny was determined by maximum likelihood method (1000 bootstrap replicates). The heat map shows fold change in leaf transcript expression one-hour post-treatment with SA, one hour post-treatment with flg22, and six hours post-infiltration with *Pseudomonas syringae* pv. *tomato* strain *hrcC*. The bar graph represents the *SnRK* expression in leaf tissue. Data were obtained from BAR Expression Angler. Blue dots represent the *SnRK* members that were cloned for yeast-two hybrid. hrcC = *P. syringae* pv. *tomato* DC3000 *hrcC* mutant defective in Type III secretion system. (B, C) Yeast two-hybrid assay of AvrPtoB and SnRK-CDPKs. AvrPtoB was co-expressed with Pto, SnRK3.1, SnRK1.1, SnRK2.6, SnRK2.8, as well as a C-terminal deletion of CPK4 (CPK4∆C) and CPK5 (CPK5∆C) in yeast. Empty vector (EV) and AD-mCherry were included as negative controls. Colony growth on SD-Leu/-Trp media confirms the presence of AD and BD vectors. Growth on SD-Leu/-Trp/-His media indicates protein-protein interaction. AD = activation domain vector, BD = binding domain vector. (D, E) Co-immunoprecipitation (Co-IP) of AvrPtoB-GFP with SnRK/CPK∆C kinase dead (KD)-HA variants in *N. benthamiana*. *AvrPtoB-GFP* under control of a dexamethasone (Dex)-inducible promoter was co-expressed with *35S::SnRKs-KD-HA* and *35S::CPKs∆C-KD-HA* in *N. benthamiana* using Agrobacterium-mediated transient expression. Expression of *AvrPtoB-GFP* was induced by 15 μM DEX for three hours at 24 hours post-infiltration. Protein extracts were subjected to anti-GFP immunoprecipitation (IP). The IP and input proteins were immunoblotted with anti-HA and anti-GFP antibodies.

Next, we analyzed the ability of diverse SnRK/CDPK members to interact with AvrPtoB by yeast two-hybrid (Supplemental Figure 2). The plant protein kinase Pto was used as a positive control (Figure 1B). SnRK1.1, SnRK2.6, and SnRK2.8 specifically interacted with AvrPtoB (Figure 1B and C). We also analyzed these CDPK members involved in flg22 signaling and NLR immune responses (Boudsocq et al., 2010; Gao et al., 2013). Auto-active CDPKs were generated by deleting their C-terminal Ca^2+^ regulatory and auto-inhibitory domains (CPKs∆C) for yeast-two hybrid screening (Klimecka and Muszyńska, 2007). CPK4∆C and CPK5∆C specifically interacted with AvrPtoB (Figure 1B, C and Supplemental Figure 2x). The expression of AvrPtoB, SnRKs, and CDPKs in yeast were verified by western blotting (Supplemental Figure 3). AvrPtoB, SnRKs, and CPKs∆C did not induce auto-activity in yeast (Figure 1C).

To validate the association of AvrPtoB with the SnRK-CDPK members identified by yeast-two hybrid, we performed co-immunoprecipitation (Co-IP) in *Nicotiana benthamiana*. To enhance the transient interaction between kinase and substrate, we employed substrate-tapping strategy with kinase-dead (KD) variants (Blanchetot et al., 2005). KD variants of five SnRK/CPK∆C members were generated by mutating the lysine residue (K) in their ATP binding pocket to alanine (A). *AvrPtoB-GFP* under the control of a dexamethasone (Dex)-inducible promoter was co-expressed with each *35S:: SnRKs-KD-HA/CPKs∆C-KD-HA* in *N. benthamiana* and immunoprecipitation (IP) was performed. Immunoblotting results demonstrated that SnRK1.1, SnRK2.6, CPK4∆CA, and CPK5∆C weakly associated with AvrPtoB (Figure 1D and E). Only SnRK2.8 strongly associated with AvrPtoB (Figure 1D and E). The strong association of SnRK2.8 with AvrPtoB in yeast and *in planta* suggests that SnRK2.8 may play an important role in AvrPtoB phosphorylation.

### AvrPtoB is phosphorylated by SnRK2.8

Next, we tested the phosphorylation of AvrPtoB by SnRK2.8 *in vitro*. Recombinant GST-AvrPtoB and GST-SnRK2.8 proteins were purified from *E. coli*, subjected to kinase activity assays, and protein phosphorylation was detected by immunoblot with antibodies recognizing pSer/pThr residues. Purified SnRK2.8 is an active kinase and exhibits strong auto-phosphorylation (Figure 2A). After incubation with SnRK2.8, AvrPtoB phosphorylation was detected in SnRK2.8 dose-dependent manner (Figure 2A). These results indicate that SnRK2.8 is able to phosphorylate AvrPtoB *in vitro*.

**Figure 2.**
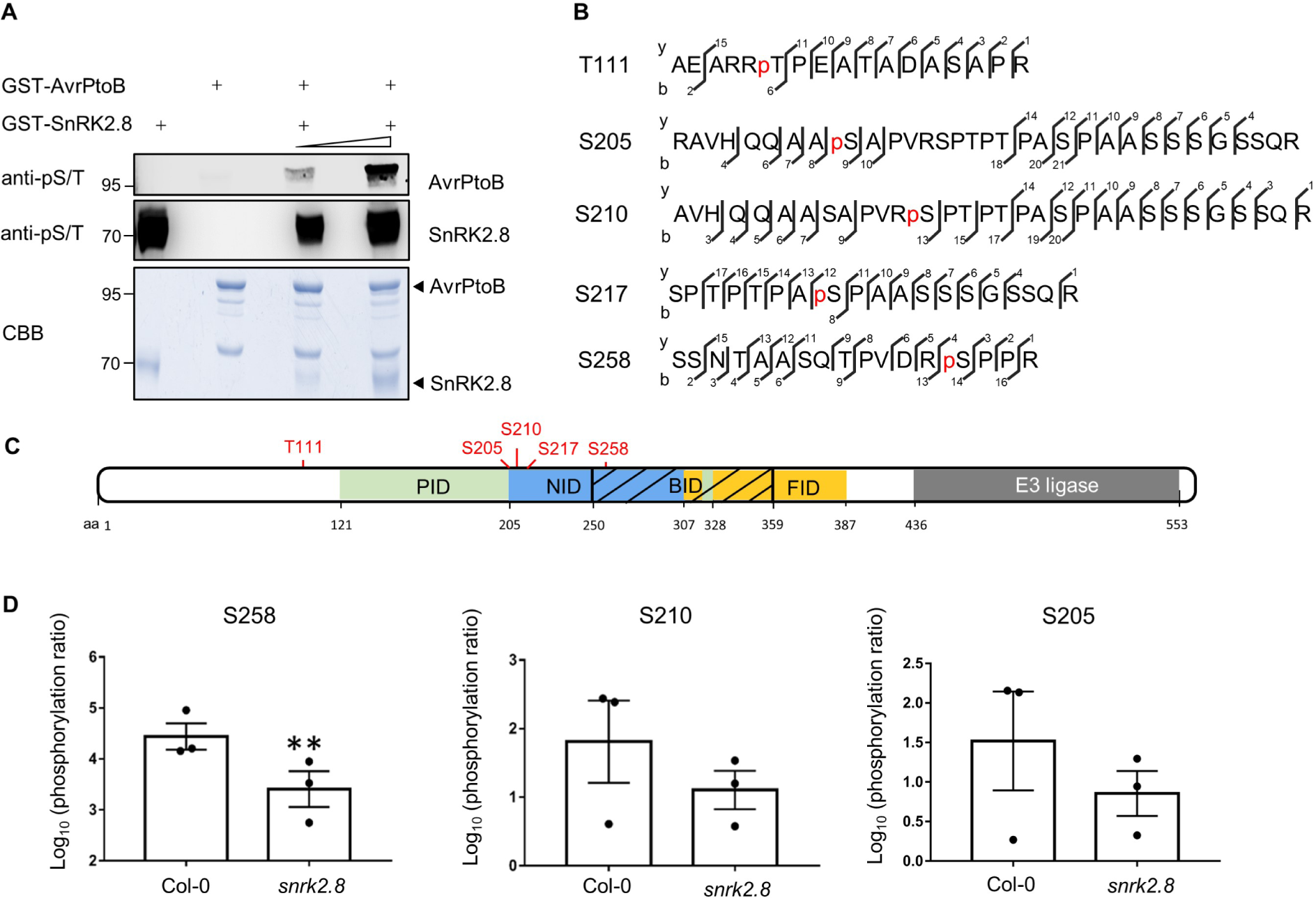
AvrPtoB is phosphorylated by SnRK2.8. (A) *In vitro* kinase activity assay of AvrPtoB and SnRK2.8. Recombinant GST-AvrPtoB and GST-SnRK2.8 proteins were purified from *E. coli* and incubated with ATP in kinase buffer for 30 min. Phosphorylation was detected by anti-pSer/Thr (anti-pS/T) immunoblot. Coomassie Brilliant Blue (CBB) staining shows protein loading. (B) AvrPtoB phosphorylation sites identified *in vivo* by LC-MS/MS. AvrPtoB-GFP was transiently expressed in *Arabidopsis* Col-0 protoplasts and total proteins were subjected to anti-GFP IP followed by trypsin digestion. Phosphorylated peptides were detected by LC-MS/MS. The observed y and b ions are numbered. (C) Diagram of phosphorylation sites and host protein interaction domains in AvrPtoB. PID (green) indicates the Pto interaction domain, NID (blue) indicates the NPR1 interaction domain, BID (black slash) indicates the BAK1 interaction domain and FID (yellow) indicates the Fen interaction domain. Numbers correspond to amino acid (aa) residues. (D) Quantification of S258, S210 and S205 phosphorylation. AvrPtoB-GFP was expressed in Col-0 and *snrk2.8* protoplasts and total proteins were subjected to anti-GFP IP followed by tryptic digestion. Phosphorylated peptides were detected by LC-MS/MS with the parallel reaction monitoring method. Peptide phosphorylation ratios (phosphorylated/non-phosphorylated) were determined using Skyline software. Data are means ± SE of three biological replicates (separate transfections). Asterisks indicate significant differences (Student’s t-test, **p<0.01).

Then, we investigated AvrPtoB phosphorylation *in vivo*. AvrPtoB-GFP was expressed in *Arabidopsis* Col-0 protoplasts, enriched by anti-GFP IP, and samples were subjected to trypsin digestion and mass spectrometry analyses (LC-MS/MS). Five phosphorylated AvrPtoB residues (T111, S205, S210, S217, and S258) were identified (Figure 2B). Previously, AvrPtoB was demonstrated to be phosphorylated at S258 in tomato, indicating that phosphorylation of this residue is conserved across diverse plant species (Xiao et al., 2007b). Aside from T111, all other phosphorylated residues map to specific domains required for binding the target proteins NPR1 (S205, S210, S217, and S258) and BAK1 (S258) (Figure 2C). The region surrounding S258 matches the proposed consensus phosphorylation motif of the SnRKs/CDPKs (Supplemental Table 1).

To determine the importance of SnRK2.8 for AvrPtoB phosphorylation *in planta*, AvrPtoB-GFP was expressed in *Arabidopsis* protoplasts from both wild-type Col-0 and the *snrk2.8* knockout. AvrPtoB-GFP was then immunoprecipitated after transfection, subjected to trypsin digestion and LC-MS/MS using parallel reaction monitoring (PRM). Using PRM, we were able to detect phosphorylation of the five previously identified AvrPtoB residues. The phosphorylation ratios (phosphorylated/nonphosphorylated peptides) of T111 and S217 were similar between Col-0 and *snrk2.8* (Supplemental Figure 4A and B, Supplemental Table 2). However, phosphorylation of S258 was significantly reduced by over 90% in *snrk2.8* and phosphorylation of S205 and S210 were reduced by~50% (Figure 2D). While the phosphorylation of S205 and S210 was decreased in the *snrk2.8* background, the decrease was not statistically significant (Figure 2D). Notably, S205, S210, and S258 are highly conserved as either S or T residues across 27 *Pseudomonas* AvrPtoB homologs (Supplemental Figure 5).

These results indicate that SnRK2.8 is one of the main kinases involved in AvrPtoB phosphorylation. However, since some phosphorylation of S205/S210/S258 remain in the *snrk2.8* knockout and *SnRK2.8* does not influence phosphorylation of T111 and S217 *in planta*, other kinases are also involved in phosphorylating AvrPtoB. AvrPtoB weakly associates with SnRK1.1, SnRK2.6 and another two CDPK members in *planta*. These kinases may also be capable of AvrPtoB phosphorylation.

### *SnRK2.8* is required for AvrPtoB virulence

AvrPtoB’s phosphorylated residues are located within regions known to bind to plant targets, which led us to investigate whether *SnRK2.8* is required for AvrPtoB virulence (Figure 2C). Since AvrPto and AvrPtoB are redundant with respect to virulence, a *Pseudomonas syringae* pv. *tomato* DC3000 *avrPto* and *avrPtoB* double knockout strain was used (Lin et al., 2005). DC3000Δ*avrPto*Δ*avrPtoB* (DC3000 −/−) was transformed with a plasmid expressing *AvrPtoB* or the empty vector (EV) control. Col-0 and *snrk2.8* plants were inoculated with DC3000 and DC3000 −/− strain variants by syringe infiltration. Four days post-inoculation DC3000 caused severe disease symptoms, while DC3000 −/− EV caused weak disease symptoms on both Col-0 and *snrk2.8* (Figure 3A). DC3000 −/− carrying *AvrPtoB* caused much more severe symptoms and higher bacterial titers compared to DC3000 −/− carrying EV on Col-0, but not the *snrk2.8* knockout (Figure 3A and B). These results indicate that *SnRK2.8* is required for AvrPtoB virulence in *Arabidopsis*.

**Figure 3.**
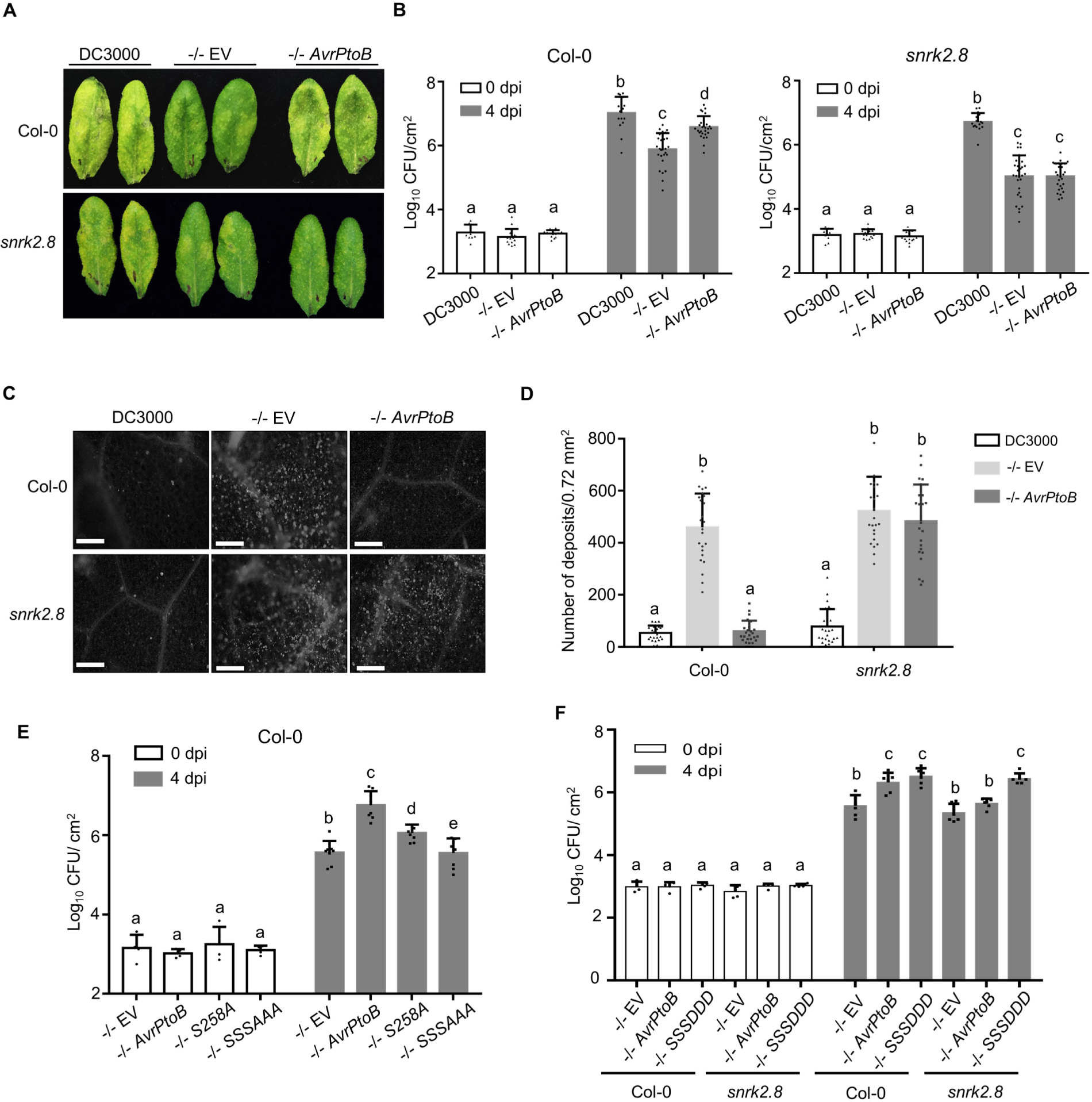
SnRK2.8 and phosphorylated serine residues are required for AvrPtoB virulence. (A) Disease symptoms of *Arabidopsis* Col-0 and the *snrk2.8* knockout after infection with *P. syringae* pv. *tomato* DC3000 (DC3000) variants. DC3000 and DC3000Δ*avrPto*Δ*avrPtoB* (−/−) carrying empty vector (EV) or expressing *AvrPtoB* on a plasmid were syringe infiltrated into *Arabidopsis* Col-0 and *snrk2.8* at a concentration of 2 × 10^5^ CFU mL^-1^. Disease symptoms were observed 4 days post-inoculation (dpi). (B) Bacterial populations in Col-0 and *snrk2.8* leaves 4 dpi. Bacterial inoculations were performed as described in (A). Log_10_ CFU/cm^2^ = log_10_ colony forming units per cm^2^ of leaf tissue. Data are means ± SD (n ≥ 9 plants for day 0, n ≥ 18 plants for day 4). Different letters indicate significant differences (two-way ANOVA, Tukey’s test, p < 0.05). (C) Callose deposition in *Arabidopsis* Col-0 and *snrk2.8* after inoculation with DC3000, −/− EV, and −/−*AvrPtoB*. Leaves were inoculated with *P. syringae* at a concentration of 1 × 10^8^ CFU mL^-1^ and harvested 16h later. Leaves were stained by 1% aniline blue and imaged by fluorescence microscopy. Scale bar, 100 μm. (D) Quantification of callose deposits. Data from three independent experiments were used for statistical analysis. Data are means ± SD (n = 24 images from 24 leaves). Different letters indicate significant differences (two-way ANOVA, Tukey’s test, p < 0.05). (E) Bacterial populations in Col-0 leaves 4 dpi. Bacterial inoculations were performed as described in (A). Log10 CFU/cm^2^ = log_10_ colony-forming units per cm^2^ of leaf tissue. Data are means ± SD (n = 4 plants for day 0, n = 8 plants for day 4). Different letters indicate significant differences (two-way ANOVA, Tukey’s test, p < 0.05). (F) Bacterial populations in Col-0 leaves 4 dpi. Bacterial inoculations were performed as described in (A). Log10 CFU/cm^2^ = log10 colony-forming units per cm^2^ of leaf tissue. Data are means ± SD (n = 4 plants for day 0, n = 6 plants for day 4). Different letters indicate significant differences (two-way ANOVA, Tukey’s test, p < 0.05).

AvrPtoB suppresses plant defenses triggered by PAMP recognition including the deposition of callose, a β-1–3 linked glucan polymer, which acts as a structural barrier (Torres et al., 2006). To investigate the impact of *SnRK2.8* on AvrPtoB virulence, we examined callose deposition. DC3000 −/− carrying EV elicited strong callose deposition in both Col-0 and *snrk2.8* (Figure 3C and D). As expected, DC3000 −/− carrying *AvrPtoB* did not elicit strong callose deposition in Col-0. However, *AvrPtoB* was unable to suppress callose deposition in the *snrk2.8* knockout (Figure 3C and D). These data indicate that AvrPtoB requires the plant kinase *SnRK2.8* for suppressing callose deposition in *Arabidopsis*.

### Phosphorylated AvrPtoB serine residues are required for virulence

To test the functional relevance of AvrPtoB phosphorylation, we analyzed the virulence of AvrPtoB phospho-null mutants. We focused on S205, S210 and S258 due to their decrease in phosphorylation in the *snrk2.8* knockout (Figure 2D). These serine residues were replaced by alanine to create single and triple phospho-null mutations. Previous research demonstrated that the alanine substitution of S258 resulted in a loss of virulence of AvrPtoB1–307 on the susceptible tomato genotype Rio Grande 76S (lacking the resistance gene Prf) (Xiao et al., 2007b). We first tested the ability of full-length AvrPtoB_S258A_ to promote bacterial growth on tomato 76S and *Arabidopsis* Col-0. The DC3000 −/− strain expressing *AvrPtoB_S258A_* was able to increase bacterial growth in both tomato and *Arabidopsis*, but not to the same extent as wild-type *AvrPtoB* (Supplemental figure 6A and B). In contrast, the DC3000 −/− strain expressing the 1-307 truncation of *AvrPtoB_S258A_* was unable to increase bacterial growth in both tomato and *Arabidopsis* (Supplemental figure 6A and B). These results indicate that the phosphorylation of S258 is important for the full virulence of AvrPtoB.

Next, we tested the virulence of the phospho-null AvrPtoB. *Arabidopsis* Col-0 was syringe infiltrated with DC3000 −/− carrying EV, *AvrPtoB, AvrPtoB*_*S258A*_, and *AvrPtoB*_*S205AS210A258A*_. As expected, infection with DC3000 −/− carrying *AvrPtoB* resulted in foliar disease symptoms, increased bacterial growth, and suppression of callose deposition compared to EV (Supplemental figure 7A, C and E). Infection with DC3000 −/− *AvrPtoB*_*S258A*_ resulted in an intermediate phenotype, with slight disease symptoms, a partial increase in bacterial growth compared to EV, and partial suppression of callose deposition compared to EV (Supplemental figure 7A, C and E). However, DC3000 −/− carrying *AvrPtoB*_*S205AS210A258A*_ was unable to induce disease symptoms, grew to the same level as the EV control, and was unable to suppress callose deposition (Supplemental figure 7A, C and E). Western blot analyses demonstrated that all AvrPtoB variants were equally expressed in DC3000 −/− (Supplemental figure 6C).

In order to investigate the importance of phosphorylation for promoting bacterial virulence, we examined bacterial growth and callose deposition in Col-0 and the *snrk2.8* knockout after infection with DC3000 −/− *AvrPtoB*_*S205DS210DS258D*_. The AvrPtoB triple phosphomimic, but not wild-type AvrPtoB, was able to enhance bacterial growth and suppress callose deposition in the *snrk2.8* knockout (Figure 3F, Supplemental Figure 7B, D and F). Collectively, these results further support the conclusion that phosphorylation of AvrPtoB by SnRK2.8 is required for effector virulence.

### AvrPtoB-mediated inhibition of NPR1 accumulation requires the plant kinase SnRK2.8

AvrPtoB associates with and ubiquitinates NPR1 in the presence of SA (Chen et al., 2017). To further investigate the relevance of SnRK2.8 for promoting AvrPtoB virulence, we examined AvrPtoB-mediated inhibition of NPR1 accumulation in Col-0 and *snrk2.8*. Both Col-0 and *snrk2.8* were pretreated with 0.5mM SA for 6 hours and then syringe inoculated with DC3000 −/− EV and DC3000 −/− *AvrPtoB*. NPR1 accumulation was detected by anti-NPR1 immunoblotting. Infection with DC3000 −/− EV induced high accumulation of NPR1 in both Col-0 and *snrk2.8* (Supplemental figure 8A and B). However, infection with DC3000 −/− *AvrPtoB* inhibited NPR1 accumulation in Col-0, but the inhibition was attenuated in the *snrk2.8* knockout, indicating that *SnRK2.8* is required for AvrPtoB-mediated inhibition of NPR1 accumulation (Supplemental figure 8A and B).

The importance of AvrPtoB phosphorylated residues for inhibition of NPR1 accumulation was examined. We co-expressed *35S::NPR1-HA* and DEX inducible *AvrPtoB-FLAG, AvrPtoB* phospho-null, and *AvrPtoB* phospho-mimetic mutants in *N. benthamiana*. Wild-type AvrPtoB, AvrPtoBS258D, and AvrPtoB_S205DS210DS258D_ significantly decreased NPR1 accumulation compared to the EV control, indicating that mimicking phosphorylation promotes virulence of AvrPtoB (Supplemental figure 8C and D). AvrPtoB_S258A_ was still able to suppress the accumulation of NPR1, indicating that the phosphorylation of serine 258 is not sufficient for this function (Supplemental figure 8C and D). In contrast, AvrPtoB_S205AS210A258A_ was unable to suppress NPR1 accumulation (Supplemental figure 8C and D). Collectively, these results show that AvrPtoB-mediated inhibition of NPR1 accumulation requires the plant kinase SnRK2.8 and the SnRK2.8-mediated phosphorylated residues.

### AvrPtoB-mediated inhibition of FLS2 accumulation and FLS2-BAK1 complex formation requires the plant kinase SnRK2.8

The flagellin receptor FLS2 is another host target of and is ubiquitinated by AvrPtoB (Gómez-Gómez and Boller, 2000; Lu et al., 2011; Göhre et al., 2008). To investigate the relevance of SnRK2.8 with AvrPtoB-mediated inhibition of FLS2 accumulation, we compared FLS2 accumulation after infection with DC3000 −/− carrying EV or *AvrPtoB* in different genetic backgrounds. Col-0 and the *snrk2.8* knockout were syringe inoculated and FLS2 accumulation was detected by anti-FLS2 immunoblotting. FLS2 accumulation was enhanced after infection with DC3000 −/− EV in both Col-0 and *snrk2.8* (Figure 4A and B). In contrast, *AvrPtoB* was able to suppress FLS2 accumulation in Col-0 but not in the *snrk2.8* knockout, demonstrating *SnRK2.8* is required for AvrPtoB-mediated suppression of FLS2 accumulation (Figure 4A and B).

**Figure 4.**
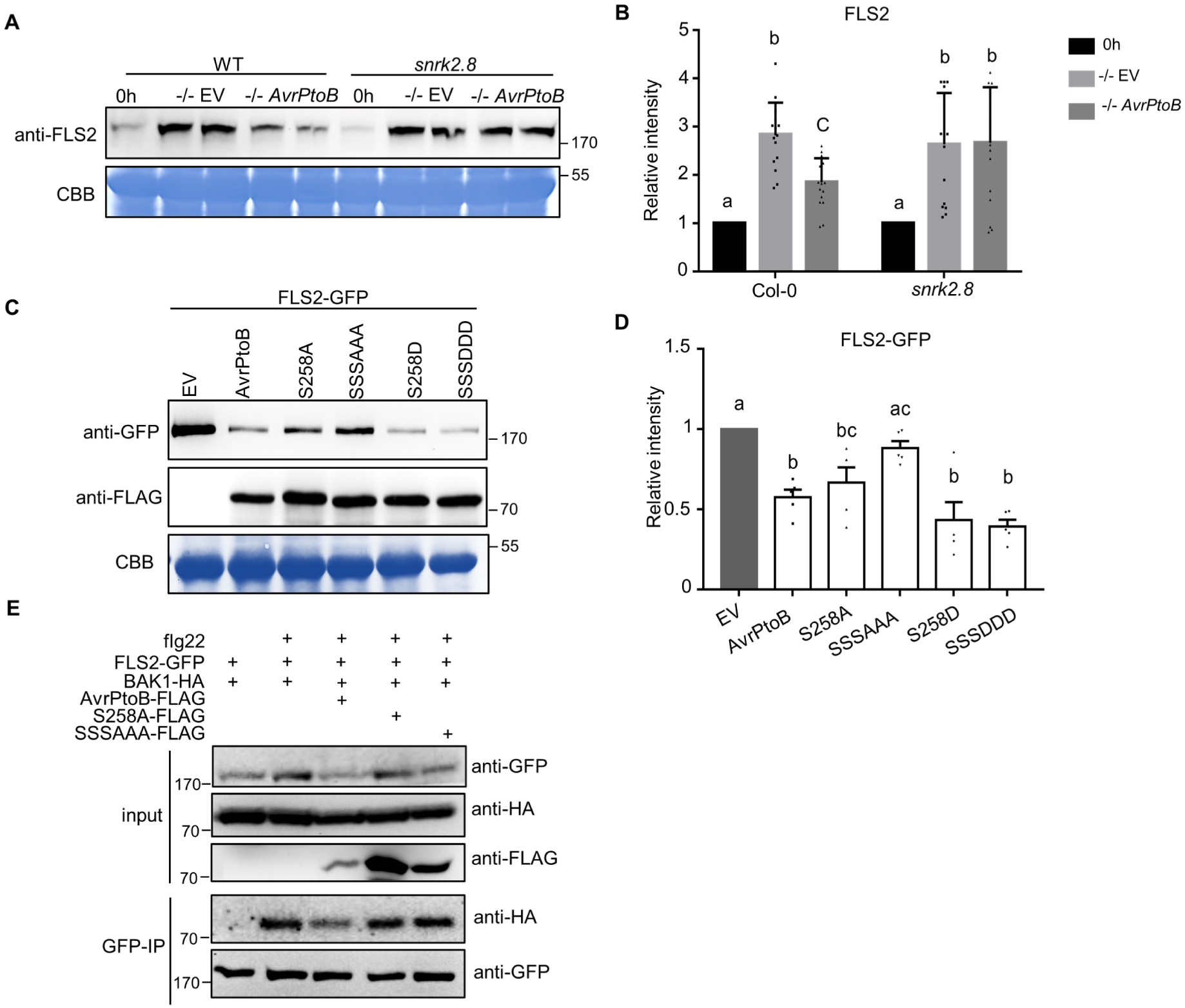
AvrPtoB-mediated degradation of FLS2 and inhibition of FLS2 complex formation requires the plant kinase SnrK2.8 and phosphorylated residues. (A) FLS2 accumulation in *Arabidopsis* Col-0 and the *snrk2.8* knockout after inoculation with *P. syringae* pv. *tomato* DC3000Δ*avrPto*Δ*avrPtoB* (−/−) variants. DC3000 −/− carrying the empty vector (EV) or a plasmid expressing wild-type *AvrPtoB* were syringe infiltrated into Col-0 and *snrk2.8* at a concentration of 1 × 10^8^ CFU mL^-1^ and proteins extracted after 8h. Protein extracts were subjected to anti-FLS2 immunoblotting. Coomassie Brilliant Blue (CBB) staining shows equal protein loading. (B) Quantification of FLS2 band intensity in (A). FLS2 band intensities were quantified by Image Lab 6.0.1. The values were normalized first by Rubisco bands and subsequently by the intensities of “0h” bands. Data are means ± SD (n = 14 leaves). Different letters indicate significant differences (one-way ANOVA, Tukey’s test, p < 0.05). (C) FLS2 accumulation in the presence of AvrPtoB phosphorylation mutants in *N. benthamiana* after Agrobacterium-mediated transient expression. *35S::FLS2-GFP* was co-expressed with FLAG-tagged Dex-inducible *AvrPtoB*, *AvrPtoB*_*S258A*_, *AvrPtoB_S205AS210AS258A_* (*SSSAAA*), *AvrPtoBS258D*, or *AvrPtoBS205DS210DS258D* (*SSSDDD*). The Dex inducible EV was used as a control. The expression of AvrPtoB-FLAG phosphorylation variants was induced by 15 μM DEX for 5 hours 24h post-Agrobacterium infiltration. Protein extracts were subjected to anti-GFP and anti-FLAG immunoblotting. CBB staining shows protein loading. (D) Quantification of FLS2-GFP band intensity in (C). FLS2-GFP bands intensities were quantified by Image Lab 6.0.1. The values were normalized first by Rubisco bands and subsequently by the intensities of “EV” treatment. The data from two independent experiments were used for statistical analysis. Data are means ± SD (n = 6 leaves). Different letters indicate significant differences (one-way ANOVA, Tukey’s test, p < 0.05). (E) Analyses of AvrPtoB-mediated inhibition of FLS2 complex formation. AvrPtoB-FLAG variants, BAK1-HA, and FLS2-GFP were co-expressed in the indicated combinations in *N. benthamiana* using Agrobacterium-mediated transient expression. The expression of AvrPtoB-FLAG phosphorylation variants was induced by 15 μM DEX for 3 hours 24h post-Agrobacterium infiltration. Leaves were infiltrated with 10 μM flg22 peptide 15 minutes before harvesting protein extracts for GFP immunoprecipitation and western blotting. Protein inputs and immunoprecipitated samples were detected by immunoblot.

To test if AvrPtoB phosphorylated residues are required for inhibition of FLS2 accumulation, we co-expressed *35S::FLS2-GFP* and DEX inducible *AvrPtoB-FLAG*, *AvrPtoB* phospho-null, and *AvrPtoB* phospho-mimetic mutants in *N. benthamiana*. Wild-type AvrPtoB, AvrPtoB_S258D_, and AvrPtoB_S205DS210DS258D_ significantly decreased FLS2 accumulation compared to the EV control (Figure 4C and D). AvrPtoB_S258A_ was still able to decrease the accumulation of FLS2, indicating that the phosphorylation of serine 258 is not sufficient for this AvrPtoB function (Figure 4C and D). In contrast, the phospho-null mutant AvrPtoB_S205AS210AS258A_ was unable to suppress FLS2 accumulation (Figure 4C and D). Collectively, these results demonstrate that AvrPtoB phosphorylated residues and the plant kinase SnRK2.8 are required for inhibition of FLS2 accumulation.

Phosphorylation can impact protein-protein interactions by creating an intricate binding surface (Tarrant and Cole, 2009). FLS2-mediated recognition of flg22 induces the formation of an FLS2-BAK1 complex, which is required for downstream immune signaling (Chinchilla et al., 2007; Shan et al., 2008). We examined flg22-induced FLS2-BAK1 association in the presence of AvrPtoB and AvrPtoB phospho-null mutants after transient expression in *N. benthamiana*. As shown in Figure 4E, wild-type AvrPtoB effectively diminished FLS2-BAK1 association in the presence of flg22, but AvrPtoB_S258A_ and AvrPtoB_S205AS210AS258A_ were unable to suppress FLS2-BAK1 complex formation

In our study, the phosphorylation of AvrPtoB is required for the effector-mediated inhibition of NPR1/FLS2 accumulation and FLS2-BAK1 complex formation. The phosphorylation of AvrPtoB may be required for the interactions with its targets or positioning of AvrPtoB’s E3 ligase domain in the appropriate orientation. S258 is in both the NPR1 and BAK1 binding domains of AvrPtoB, while S205 and S210 are present in the NPR1 binding domain. Though the single phospho-null S258A mutant was able to inhibit NPR1/FLS2 accumulation, it was unable to disrupt FLS2-BAK1 complex formation. However, the triple phospho-null mutant completely lost both of these functions, indicating that the phosphorylation of S205 and/or S210 is required for AvrPtoB-mediated target degradation (Figure 4C, D and Supplemental figure 8C, D). These data are consistent with previous observations that AvrPtoB is a multifunctional protein with different interfaces involved in interacting with diverse host proteins (Martin, 2011).

Despite the importance of effector phosphorylation, the plant kinases involved remain largely elusive. Here, we identified a member of the SnRK-CDPK family, SnRK2.8, whose phosphorylation of the conserved bacterial effector AvrPtoB promotes effector virulence activities (Supplemental figure 9). The SnRK-CDPK family is conserved in plants and most reported effectors phosphorylated residues are consistent with SnRK-CDPK phosphorylation site preferences. Thus, we hypothesize that different SnRKs and CPDKs may be involved in phosphorylation of diverse effector proteins. An increasing number of studies show the key roles of SnRKs in plant responses to different types of pathogens. Tomato SnRK1 phosphorylates the pathogenic protein βC1 of the tomato yellow leaf curl virus to limit viral infection and is also involved in cell death elicited by *Xanthomonas* effectors (Shen et al., 2011; Szczesny et al., 2010; Avila et al., 2012). Future investigation of the role of SnRK-CDPKs in phosphorylation of diverse pathogen effectors will provide a more comprehensive understanding of how pathogen effectors exploit their hosts to facilitate disease development.

## Methods

### Plant material and growth conditions

*Arabidopsis thaliana* seeds were stratified for two days at 4℃ in the dark before sowing. Plants were grown in a controlled environmental chamber at 23℃ and 70% relative humidity with a 10-hr-light /14-hr-dark photoperiod (100 μM m^-2^ s^-1^). Four-to-five-week old plants were used for all experiments. A confirmed *snrk2.8* knockout T-DNA insertion line (SALK_073395) in the Columbia (Col-0) background was obtained from the *Arabidopsis* Biological Resource Center (Shin et al., 2007).

*N. benthamiana* was grown in a growth chamber at 26℃ with a 16-hr-light/8-hr-dark photoperiod (180 μM m^-2^ s^-1^). Three-to-four-week old plants were used for Agrobacterium-mediated transient protein expression.

### Yeast two-hybrid screen and molecular cloning

The GAL4 based Matchmaker yeast two-hybrid system was used for the AvrPtoB-kinase interaction screen (Clonetech). Cloning details are provided in supplemental data.

### Phylogeny and expression of *SnRKs*

A total of 38 *Arabidopsis* SnRK protein sequences were obtained from the *Arabidopsis* information resource (TAIR, http://Arabidopsis.org). Multiple sequence alignments were performed with MEGA X (Kumar et al., 2018). Phylogeny was determined by maximum likelihood method (1000 bootstrap replicates). *SnRK* transcriptional expression in leaves were obtained from BAR Expression Angler (Austin et al., 2016).

### Bacterial growth assay

The broad host range plasmid pBAV226 was used to express *AvrPtoB* variants. pBAV226 harboring *AvrPtoB* or its derivatives were electroporated into the *P. syringae* pv. *tomato* DC3000Δ*avrPto*Δ*avrPtoB* (DC3000 −/−) strain (Lin and Martin, 2005). Bacteria were syringe infiltrated into five-week-old *Arabidopsis* Col-0 or *snrk2.8* leaves at a concentration of 2 × 10^5^ CFU m^L-1^ (OD600 = 0.0002). Bacterial titers were determined as colony forming units (CFU) per cm^2^ at four days post inoculation as previously described (Liu et al., 2009).

### Callose staining and quantification

*Arabidopsis* plants were inoculated with *P. syringae* at a concentration of 1 × 10^8^ CFU mL^-1^ (OD600 = 0.1) for 16 hr. Infected leaves were fixed and destained in 95% ethanol, washed twice with 70% ethanol and three times with distilled water followed by staining with 1% aniline blue in 150 mM K_2_HPO_4_ (pH 9.5) for 30 min in dark at RT. Callose deposition was imaged by fluorescence microscopy (Leica CMS GmbH) using a DAPI filter and Image J (Collins, 2007).

### Statistical analyses

Statistical analyses were performed by Prism 7 software (GraphPad). The data are presented as mean ± SD or ± SE. For quantification of phosphorylated peptides and quantification of NPR1 or FLS2 band intensity, n represents the number of experimental replicates. For bacterial growth and callose deposition, n represents the number of individual plants. Student’s t test was used to compare means for two groups. One-way ANOVA or two-way ANOVA with Turkey’s multiple-comparison test was performed for multiple comparisons. Statistical analyses, p-values, and the exact value of n are described in detail in the figures and figure legends.

**Protein extraction, purification, interactions and phosphorylation analyses are included in Supplemental methods.**

## Author Contributions

G.C. conceived the study, L.L. and G.C. designed experiments and wrote the manuscript. L.L. performed most experiments. D.M.S. analyzed the AvrPtoB homologs. All authors approved of the final manuscript.

## Acknowledgements

The authors thank Sheng Luan (University of California, Berkeley) for providing clones of different *SnRK* members and Michelle Salemi (University of California, Davis) for assistance in mass spectrometry analyses.

