## Supplemental figures, tables, methods for "Identification of a plant kinase that phosphorylates the bacterial effector AvrPtoB"

**Supplemental Table and Figure Legends.**

**Supplemental Table 1. The phosphorylation sites of bacterial type III effectors identified by mass spectrometry. Related to Figure 1.** Data were collected from the references in the table. Red indicates the phosphorylated residue.

**Supplemental Table 2. Related to Figure 2 and Supplemental Figure 4.** AvrPtoB was expressed in Col-0 and *snrk2.8* protoplasts and total proteins were subjected to anti-GFP IP followed by tryptic digestion. Phosphorylated peptides were detected by LC-MS/MS with the parallel reaction monitoring method. Peptide phosphorylation ratios were determined using Skyline software. Phoshorylation intensity is reported for residue T111 as no unphosphorylated peptides were detected by MS. Data are means ± SE of three biological replicates (separate transfections).

**Supplemental Table 3. Primers used in this study. Related to Materials and Methods.**

**Supplemental Table 4. Isolation list for PRM. Related to Figure 2.** Modified and unmodified peptide sequences used to quantify phosphorylation by PRM are noted. CID = collision induced dissociation, m/z = mass to charge, z = charge state.

**Supplemental Figure 1. Amino acid conservation surrounding known effector phosphorylation sites. Related to Figure 1.**

WebLogo alignment of the effector phosphorylation sites shown in Table S1. Red arrows point the phosphorylated residues. Amino acid size correlates with degree of conservation. The sequence logo was created by the WebLogo 3 (Crooks et al., 2004).

**Supplemental Figure 2. Yeast-two hybrid screening of interactions between the *Pseudomonas syringae* effector AvrPtoB and SnRK or CDPK members. Related to Figure 1.**

Yeast two-hybrid (Y2H) assay of AvrPtoB and SnRK-CDPKs in the Matchmaker GaL4 system. AvrPtoB was expressed from the pGADT7 vector, while SnRKs and C-terminal deletions of CPKs (CPKs△C) were expressed from the pGBKT7 vector in *Saccharomyces cerevisea.* The pGBKT7 empty vector (EV) was included as a negative control. Blue colonies on SD -Leu/-Trp/-His/X-α-gal media indicate protein-protein interactions. AD = activation domain vector pGADT7, BD = binding domain vector pGBKT7.

**Supplemental Figure 3. Protein expression of *SnRKs*, *CPKs*, and *AvrPtoB* used in the Y2H assay. Related to Figure 1.**

Yeast proteins were extracted and SnRK and CPK proteins were detected by anti-myc immunoblot and AvrPtoB was detected by anti-HA immunoblot. Expression of the positive (Pto) and negative (mCherry) controls used in Figure 1 are shown.

**Supplemental Figure 4. AvrPtoB T111 and S217 phosphorylation levels in the *snrk2.8* knockout compared to wild-type Col-0. Related to Figure 2.**

AvrPtoB-GFP was expressed in Col-0 and *snrk2.8* protoplasts, total proteins were subjected to anti-GFP IP followed by tryptic digestion. Phosphorylated peptides were detected by LC-MS/MS with the parallel reaction monitoring method. The peptide phosphorylation intensity of T111 (A) and the phosphorylation ratio of S217 (B) were determined using Skyline software. Data are means ± SE of three biological replicates (separate transfections). No significant differences were detected (Student's t-test, *$p<0.05$).

**Supplemental Figure 5. Conservation of phosphorylated residues in different AvrPtoB homologs. Related to Figure 2.**

AvrPtoB phosphorylated residues identified by mass spectrometry were analyzed for their conservation across 27 AvrPtoB homologs from *P. syringae.* The translated sequence of AvrPtoB was used in BLAST searches against the NCBI nr database on January 5th, 2020 (*Pseudomonas syringae* group, taxid: 136849; default parameters otherwise) to identify homologs. Hits were filtered based on a minimum 65% sequence identity, a minimum 85% query coverage and an E-value of 0. Sequences were downloaded and aligned using Genious alignment (default parameters). The percentage of identity is shown by gradient white to black (white means less than 60% similar, gray means 60-80% similar, dark gray means 80-100%, and black means 100% similar). AvrPtoB phosphorylation sites identified *in vivo* are shown with arrows.

**Supplemental Figure 6. AvrPtoB S258 is required for virulence in tomato and *Arabidopsis*. Related to Figure 3.**

(A) Bacterial populations in the susceptible tomato genotype Rio Grande 76S five days post-inoculation with *P. syringae* pv. *tomato* DC3000Δ*avrPto*Δ*avrPtoB* (-/-) variants. DC3000 -/- carrying empty vector (EV) or plasmids expressing wild-type *AvrPtoB*, *AvrPtoB*$_{S258A}$, *AvrPtoB*$_{1-307}$, and *AvrPtoB*$_{1-307S258A}$ were dip inoculated on tomato at a concentration of $2 \times 10^8$ CFU $mL^{-1}$ with 0.005% Silwet. $Log_{10}$ CFU/$cm^2$ = $log_{10}$ colony-forming units per $cm^2$ of leaf tissue. Data are means ± SD (n ≥ 6 plants). Different letters indicate significant differences (one-way ANOVA, Tukey's test, $p < 0.05$).

(B) Bacterial populations in *Arabidopsis* Col-0 three days post-inoculation with DC3000 -/- variants. DC3000 -/- variants as described in (A) were syringe infiltrated in Col-0 at a concentration of $2 \times 10^5$ CFU $mL^{-1}$. Data are means ± SD (n = 4 plants for day 0, n = 6 plants for day 3). Different letters indicate significant differences (two-way ANOVA, Tukey's test, $p < 0.05$).

(C) Expression of AvrPtoB-HA and its derivatives from DC3000 -/- . Bacteria were grown in minimal media at 18 °C. Anti-HA western blotting was performed to detect the expression of AvrPtoB in bacterial pellets.

**Supplemental Figure 7. Phosphorylated AvrPtoB serine residues are required for virulence. Related to Figure 3.**

(A) Disease symptoms of *Arabidopsis* Col-0 after inoculation with *P. syringae* pv. *tomato* DC3000Δ*avrPto*Δ*avrPtoB* (-/-) variants. DC3000 -/- carrying empty vector (EV) or plasmids expressing wild-type *AvrPtoB*, *AvrPtoB*$_{S258A}$, or *AvrPtoB*$_{S205AS210AS258A}$ (*SSSAAA*) were syringe infiltrated into *Arabidopsis* Col-0 at a concentration of $2 \times 10^5$ CFU mL$^{-1}$. Disease symptoms were observed 4 days post-inoculation (dpi).

(B) Disease symptoms of *Arabidopsis* Col-0 after inoculation with *P. syringae* pv. *tomato* DC3000Δ*avrPto*Δ*avrPtoB* (-/-) variants. DC3000 -/- carrying empty vector (EV) or plasmids expressing wild-type *AvrPtoB*, or *AvrPtoB*$_{S205DS210DS258D}$ (*SSSDDD*). Plants were treated as described in (A).

(C) Callose deposition in *Arabidopsis* Col-0 after inoculation with -/- carrying EV, *AvrPtoB*, *S258A*, and *SSSAAA*. Leaves were inoculated with *P. syringae* at a concentration of $1 \times 10^8$ CFU mL$^{-1}$ and harvested 16h later. Leaves were stained by 1% aniline blue and imaged by fluorescence microscopy. Scale bar, 100 μm.

(D) Callose deposition in *Arabidopsis* Col-0 after inoculation with -/- carrying EV, *AvrPtoB*, and *SSSDDD*. Leaves were inoculated and callose visualized as described in (C).

(E) Quantification of callose deposits. Data are means ± SD (n = 8 images from 8 leaves for -/- EV, n = 18 images from 18 leaves for rest variants). Different letters indicate significant differences (one-way ANOVA, Tukey's test, $p < 0.05$).

(F) Quantification of callose deposits. Data are means ± SD (n = 21 images from 21 leaves for -/- EV, n = 30 images from 30 leaves for rest variants). Different letters indicate significant differences (one-way ANOVA, Tukey's test, $p < 0.05$).

**Supplemental Figure 8. AvrPtoB mediated degradation of NPR1 requires the plant kinase SnrK2.8 and phosphorylated residues. Related to Figure 4.**

(A) NPR1 accumulation in *Arabidopsis* Col-0 and the *snrk2.8* knockout after inoculation with *P. syringae* pv. *tomato* DC3000Δ*avrPto*Δ*avrPtoB* (-/-) variants. DC3000 -/- carrying the empty vector (EV) or a plasmid expressing wild-type *AvrPtoB* were syringe infiltrated into Col-0 and *snrk2.8* at a concentration of $1 \times 10^8$ CFU $mL^{-1}$ and proteins extracted after four hours. Protein extracts were subjected to anti-NPR1 immunoblotting. Coomassie Brilliant Blue (CBB) staining shows equal protein loading.

(B) Quantification of NPR1-HA band intensity in (C). NPR1-HA bands intensities were quantified by Image Lab 6.0.1 (BIO-RAD). The values were normalized first by Rubisco bands and subsequently by the intensities of "EV" treatment. Data are means ± SD (n = 6 leaves). Different letters indicate significant differences (one-way ANOVA, Tukey's test, $p < 0.05$).

(C) NPR1 accumulation in the presence of AvrPtoB phosphorylation mutants in *N. benthamiana* after Agrobacterium-mediated transient expression. *35S::NPR1-HA* was co-expressed with FLAG-tagged Dex-inducible *AvrPtoB*, $AvrPtoB_{S258A}$, $AvrPtoB_{S205AS210AS258A}$ (*SSSAAA*), $AvrPtoB_{S258D}$ or $AvrPtoB_{S205DS210DS258D}$ (*SSSDDD*). The Dex inducible EV was used as a control. The expression of AvrPtoB-FLAG phosphorylation variants was induced by 15 μM DEX for 5 hours 24h post-Agrobacterium infiltration. Protein extracts were subjected to anti-HA and anti-FLAG immunoblotting. CBB staining shows protein loading.

(D) Quantification of NPR1 band intensity in (A). NPR1 bands intensities were quantified by Image Lab 6.0.1 (BIO-RAD). The values were normalized first by Rubisco bands and subsequently by the intensities of "0h" bands. Data are means ± SD (n = 14 leaves). Different letters indicate significant differences (one-way ANOVA, Tukey's test, $p < 0.05$).

**Supplemental Figure 9. Model of AvrPtoB phosphorylation by SnRK2.8. Related to Figure 1-4 and Supplemental figure 7, 8.**

(A) SnRK2.8 phosphorylates AvrPtoB in *Arabidopsis* Col-0 and the phosphorylated AvrPtoB mediates degradation of FLS2/NPR1 and inhibition of FLS2 complex formation.

(B) AvrPtoB virulence function is lost in *snrk2.8* knockout.

### Supplemental methods

**Yeast two-hybrid screen**

The GAL4 based Matchmaker yeast two-hybrid system was used for the AvrPtoB-kinase interaction screen (Clonetech). *AvrPtoB* and a *mCherry* negative control were cloned into the pGADT7 vector fused to the GAL4 activation domain and HA tag. *Arabidopsis SnRK*s and *CDPKs* were cloned into the pGBKT7 vector, which contains the GAL4 DNA binding domain and N-terminal Myc epitope tag. In order to detect interactions with CDPKs, their C-terminal $Ca^{2+}$ regulatory and auto-inhibitory domains were removed prior to clone into the pGBKT7 vector. Primers are listed in Table S2. The pGADT7-*AvrPtoB* and pGADT7-*mCherry* plasmids were separately co-transformed with each pGBKT7-*SnRK/CPKΔC* plasmid into the yeast strain AH109, colonies were selected on SD -Leu/-Trp dropout media and tested for interactions on SD -Leu/-Trp/-His dropout media containing X-α-Gal. Yeast transformation and media preparation were performed per manufacturer instructions (Clontech). To confirm protein expression, yeast proteins were extracted as described previously and subjected to anti-HA HRP (Roche #12013819001; 1:2,000) and anti-Myc (Clontech #631206; 1: 2,000) immunoblotting (Zhang et al., 2011b).

**Plant protein extraction and immunoblotting**

Plant tissues were ground in liquid nitrogen and homogenized in Protein Extraction Buffer [(PEB: 50 mM Tris-HCl, pH 7.5, 1 mM EDTA, pH 8.0, 150 mM NaCl, 0.1% Triton X-100, 0.5% IGEPAL, 5% glycerol, 1 mM PMSF, 3 mM DTT, 1 × CPI (Thermo Fisher Scientific), 1× PPI (Thermo Fisher Scientific), 50 mM MG132 (Sigma-Aldrich)] (Chen et al., 2017). The homogenate was cleaned by centrifuging at 14,000 rpm for 15 min at 4 ℃, and boiled with 5 × SDS buffer (250 mM Tris-HCl pH 6.8, 6% SDS, 0.5 M DTT, 30% glycerol, 0.08% bromophenol blue) for 10 min.

**Co-immunoprecipitation assays**

To confirm the association between AvrPtoB and SnRK/CDPKs *in planta*, kinase dead variants were generated and tested for their ability to associate with wild-type AvrPtoB after transient expression in *N. benthamiana*. *SnRK1.1-KD* (K71A), *SnRK2.6-KD* (K50A), *SnRK2.8-KD* (K33A), *CPK4ΔC-KD* (K54A) and *CPK5ΔC-KD* (K126A) kinase dead variants were generated by PCR-based site-directed mutagenesis and fused with a C-terminal HA tag in the binary vector pGWB414 (Nakagawa et al., 2007). Primers are listed in Table S3. *AvrPtoB* fused with a C-terminal GFP tag was cloned into the dexamethasone (Dex)-inducible binary vector pTA7001 (Gu and Innes, 2011). Binary vectors were transformed into *Agrobacterium tumefaciens* GV3101. *Agrobacterium* suspensions were co-infiltrated into *N. benthamiana* leaves at an OD600 = 0.4 for AvrPtoB and an OD600 = 0.6 for each kinase. Twenty-four hours post-inoculation, 15 μM DEX and 0.01% Silwet L-77 were sprayed to induce the expression of AvrPtoB-GFP. Two grams of leaf tissue per sample were collected three hours post-DEX application.

For immunoprecipitations, *N. benthamiana* leaf tissues were ground in liquid nitrogen and re-suspended in 2 mL IP buffer (50mM Tris-HCL ph7.5, 150mM NaCl, 0.1% Triton, 0.2% NP-40, 1× complete protease inhibitor (Thermo Fisher Scientific #A32963) and 1× phosphatase inhibitor (Thermo Fisher Scientific #A32957), 1mM DTT, 40uM MG132, 0.5% PVP). Samples were centrifuged at 14,000 rpm for 15 min and filtered to remove debris using a poly-prep chromatography column (10 mL, Bio-Rad). The supernatant was incubated with 25 μL of anti-GFP agarose beads at 4°C for 1.5 hr. Beads were washed once with IP buffer by centrifuging at 3000 rpm for two min and twice by filtration through a pierce centrifuge column (0.8 mL, Bio-Rad). Proteins were eluted from the beads by boiling in 2× Laemmli buffer for five min. Proteins were separated by SDS-PAGE and immunoblotted with anti-HA HRP (Roche #12013819001; 1:2,000) and anti-GFP HRP (Miltenyi Biotec #130-091-833; 1:3,000).

To test the ability of AvrPtoB phosphorylation mutants to disrupt FLS2-BAK1 complex formation, *AvrPtoB* phospho-null mutants (S258A and S205AS210AS258A) were generated by PCR-based site-directed mutagenesis. Primers are listed in Table S2. *AvrPtoB* variants were fused with C-terminal 3× FLAG tag in binary vector pTA7001, and *FLS2* and *BAK1* were separately fused with C-terminal GFP tag and HA tag in binary vector pEarleyGate 103 and pGWB14 (Earley et al., 2006; Nakagawa et al., 2007). The binary vectors were transformed into *Agrobacterium tumefaciens* GV3101. *FLS2*, *BAK1*, and *AvrPtoB* variants *Agrobacterium* suspensions were co-infiltrated into *N. benthamiana* leaves, 15 μM DEX and 0.01% Silwet L-77 were sprayed to induce the expression of AvrPtoB at 24 hpi for three hours, two grams of leaf tissue for each sample were collected after 15 min treatment with 5 mM $MgCl_2$ or 10 μM flg22. Immunoprecipitation was performed as described above.

**Recombinant protein purification**

*AvrPtoB* and *SnRK2.8* were cloned in *E.coli* expression vector pDEST15 (Invitrogen) fused with N-terminal GST tag, the constructs were transformed into *E.coli* BL21 (DE3). 200 mL of *E.coli* culture was grown at 28 ℃ until OD600 = 0.5. Protein expression was induced with 0.5mM IPTG at 16 ℃ for 12 hr. Cells were harvested by centrifuging at 5000 g 4 ℃ for 10 min and washed once with buffer A (0.1 M Tris-HCl pH 7.5, 150 mM NaCl, 1 mM PMSF, 1 × CPI, 10 mM DTT and 10 uM MG132). Cell pellets were resuspended in 3 mL of buffer A with 15 μg/mL lysozyme and incubated on ice for 30 min. Total protein was released by sonication and incubated with Glutathione Sepharose 4B (GE Healthcare #GE17-0756-01) at 4 ℃ for one hour. Agarose beads were washed three times with buffer A by centrifuging at 5000 g for 5 min. Proteins were eluted by incubating with buffer B (50 mM Tris-HCl pH8, 10 mM reduced Glutathione) for 10 min at room temperature (RT).

**Kinase activity assay**

An *in vitro* kinase activity assay was performed with recombinant proteins, 3 μg of GST-AvrPtoB and 0.3-1 μg GST-SnRK2.8 were mixed in kinase buffer (20 mM Tris-HCl pH7.5, 10 mM $MgCl_2$, 1 mM $CaCl_2$, 100 μM ATP, 1 mM DTT). The kinase reaction was performed at 30 ℃ for 30 min and stopped by 3 × Laemmli buffer. Protein samples were separated in SDS-PAGE and immunoblotted with anti-pSer/Thr antibody (Sigma #P3430; 1:1000) and anti-GST antibody (Sigma; 1: 3000).

**Phosphorylation site identification and quantification**

To identify AvrPtoB phosphorylation sites *in vivo*, *AvrPtoB-YFP* in the pBluescript vector was transiently expressed in Col-0 and *snrk2.8* protoplasts, protoplast preparation and transient transformation were performed as previously described (Yoo et al., 2007). 1 mL of protoplasts was transfected with 100 µg of plasmid and collected in 9 hr. Protein was released in IP buffer (without 0.5% PVP) and subject to GFP-IP as described above. Protein peptides were generated by in-solution trypsin digest as previously described and subjected to LC-MS/MS run by Orbitrap Fusion Lumos mass spectrometer (Thermo Fisher Scientific) (Minkoff et al., 2013). LC-MS/MS data was analyzed by software MaxQuant (Tyanova et al., 2016).

To quantify the phosphorylated peptides, an inclusion list of phosphopeptide and control peptides, including the Mono-isotopic precursor (m/z) and charge state (z), was generated by software Skyline based on previous MS data, as shown in Table S4 (MacLean et al., 2010). The peptide samples were scanned by Orbitrap Fusion Lumos mass spectrometer (Thermo Fisher Scientific) with a parallel reaction monitoring (PRM) method. The PRM data were analyzed by MaxQuant and Skyline, the peptides peak areas were exported for the quantification analysis.

**NPR1 and FLS2 accumulation**

To test the ability of AvrPtoB to inhibit NPR1 and FLS2 accumulation in *Arabidopsis* Col-0 and the *snrk2.8* knockout. DC3000 -/- carrying the empty vector (EV) or a plasmid expressing wild-type *AvrPtoB* were syringe infiltrated into Col-0 and *snrk2.8* at a concentration of $1 \times 10^8$ CFU $mL^{-1}$ and proteins extracted after 4h (NPR1) and 8h (FLS2). The immunoblot was performed by anti-NPR1 (Agrisera #AS12 1854; 1:1000) and anti-FLS2 (Agrisera #AS12 1857; 1:5000) primary antibody followed by anti-rabbit-HRP (BioRad #170-5046; 1:3000) secondary antibody.

To test the ability of AvrPtoB phosphorylation mutants to inhibit NPR1 and FLS2 accumulation, *AvrPtoB* phospho-null mutants (S258A and S205AS210AS258A) and *AvrPtoB* phospho-mimic mutants (S258D and S205DS210DS258D) were generated by PCR-based site-directed mutagenesis. Primers are listed in Table S2. *AvrPtoB* variants were fused with C-terminal 3× FLAG tag in binary vector pTA7001, and *NPR1* and *FLS2* were separately fused with C-terminal HA tag and GFP tag in binary vector pEarleyGate 103 and pGWB14 (Earley et al., 2006; Nakagawa et al., 2007). The immunoblot was performed by anti-HA HRP (Roche #12013819001; 1:2,000), anti-FLAG (Sigma #A8592; 1:3000), and anti-GFP HRP (Miltenyi Biotec #130-091-833; 1:3,000).

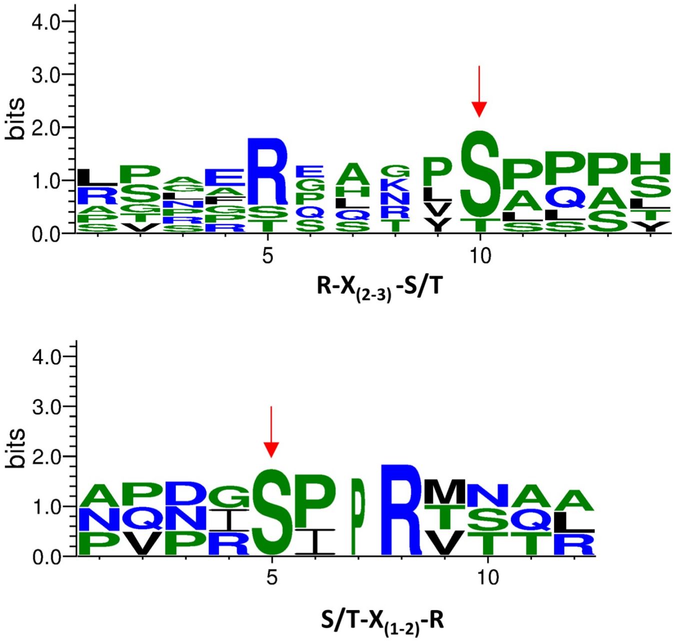


**Supplemental Figure 1. Amino acid conservation surrounding known effector phosphorylation sites. Related to Figure 1.**

WebLogo alignment of the effector phosphorylation sites shown in Table S1. Red arrows point the phosphorylated residues. Amino acid size correlates with degree of conservation. The sequence logo was created by the WebLogo 3 (Crooks et al., 2004).

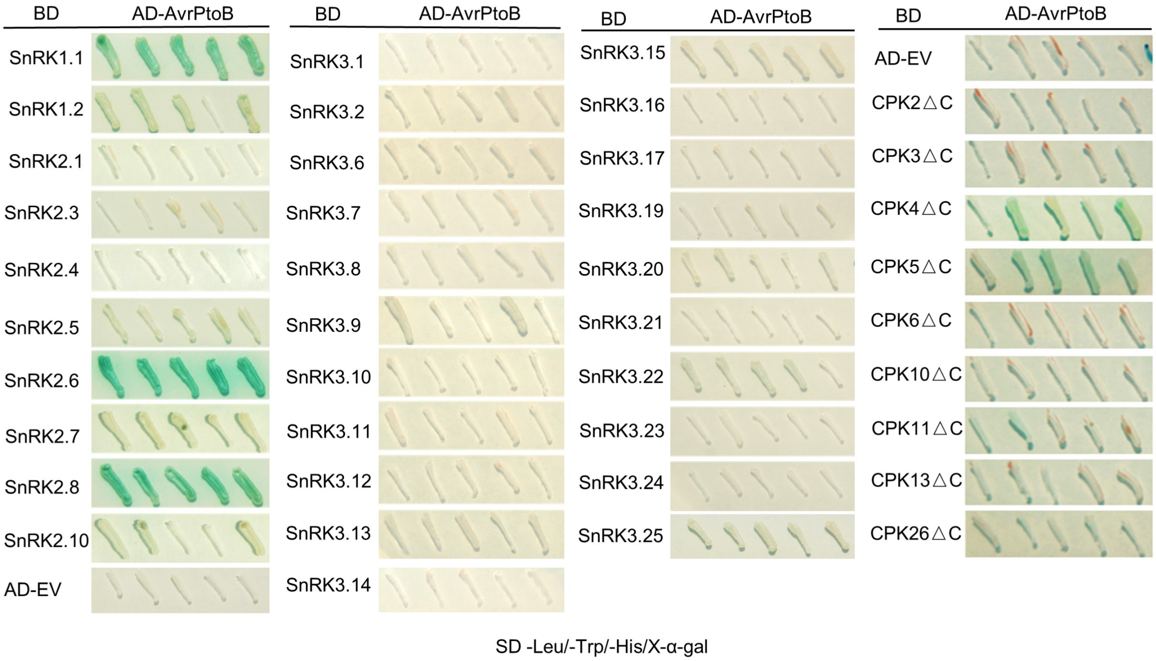


**Supplemental Figure 2. Yeast-two hybrid screening of interactions between the *Pseudomonas syringae* effector AvrPtoB and SnRK or CDPK members. Related to Figure 1.**

Yeast two-hybrid (Y2H) assay of AvrPtoB and SnRK-CDPKs in the Matchmaker GaL4 system. AvrPtoB was expressed from the pGADT7 vector, while SnRKs and C-terminal deletions of CPKs (CPKs△C) were expressed from the pGBKT7 vector in *Saccharomyces cerevisea.* The pGBKT7 empty vector (EV) was included as a negative control. Blue colonies on SD -Leu/-Trp/-His/X-α-gal media indicate protein-protein interactions. AD = activation domain vector pGADT7, BD = binding domain vector pGBKT7.

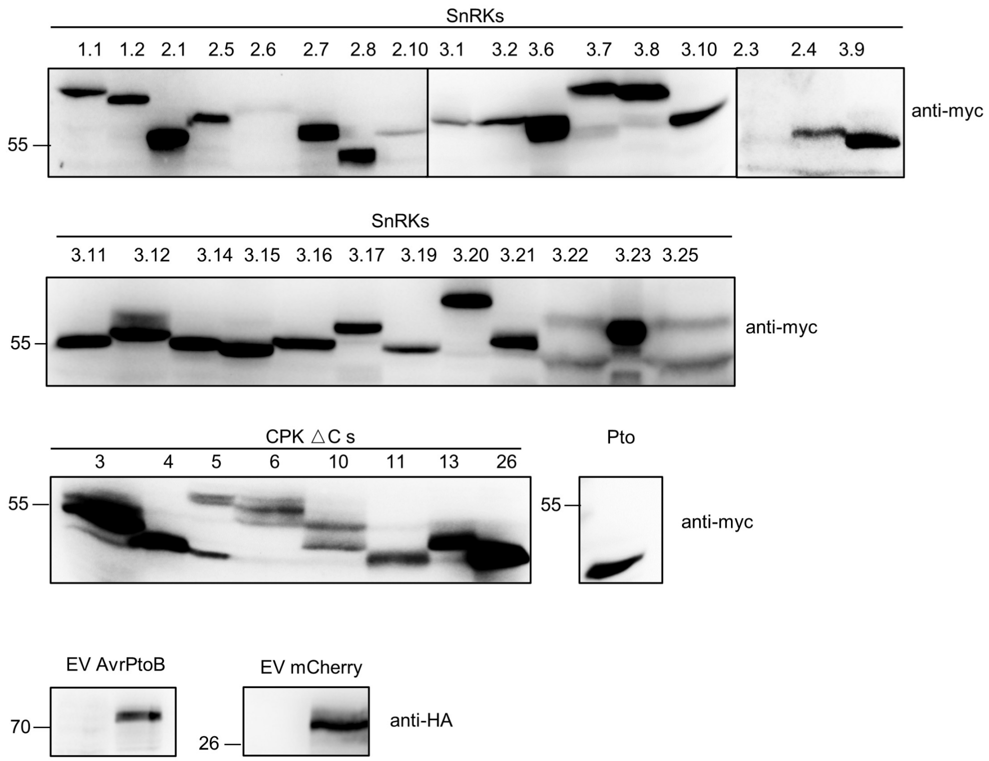


**Supplemental Figure 3. Protein expression of *SnRKs*, *CPKs*, and *AvrPtoB* used in the Y2H assay. Related to Figure 1.**

Yeast proteins were extracted and SnRK and CPK proteins were detected by anti-myc immunoblot and AvrPtoB was detected by anti-HA immunoblot. Expression of the positive (Pto) and negative (mCherry) controls used in Figure 1 are shown.

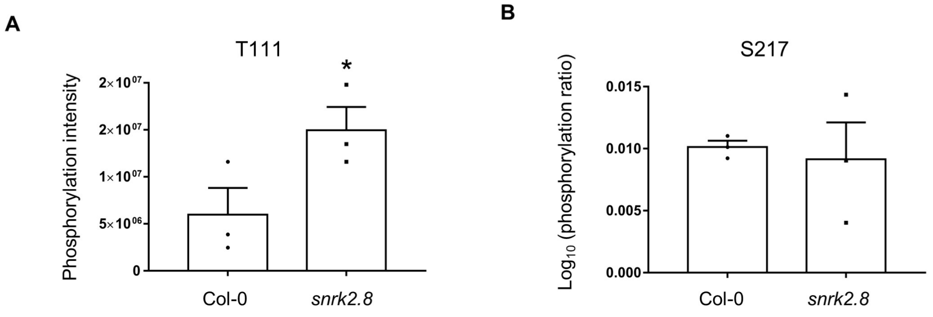


**Supplemental Figure 4. AvrPtoB T111 and S217 phosphorylation levels in the *snrk2.8* knockout compared to wild-type Col-0. Related to Figure 2.**

AvrPtoB-GFP was expressed in Col-0 and *snrk2.8* protoplasts, total proteins were subjected to anti-GFP IP followed by tryptic digestion. Phosphorylated peptides were detected by LC-MS/MS with the parallel reaction monitoring method. The peptide phosphorylation intensity of T111 (A) and the phosphorylation ratio of S217 (B) were determined using Skyline software. Data are means ± SE of three biological replicates (separate transfections). Asterisks indicate significant differences (Student's t-test, *$p<0.05$).

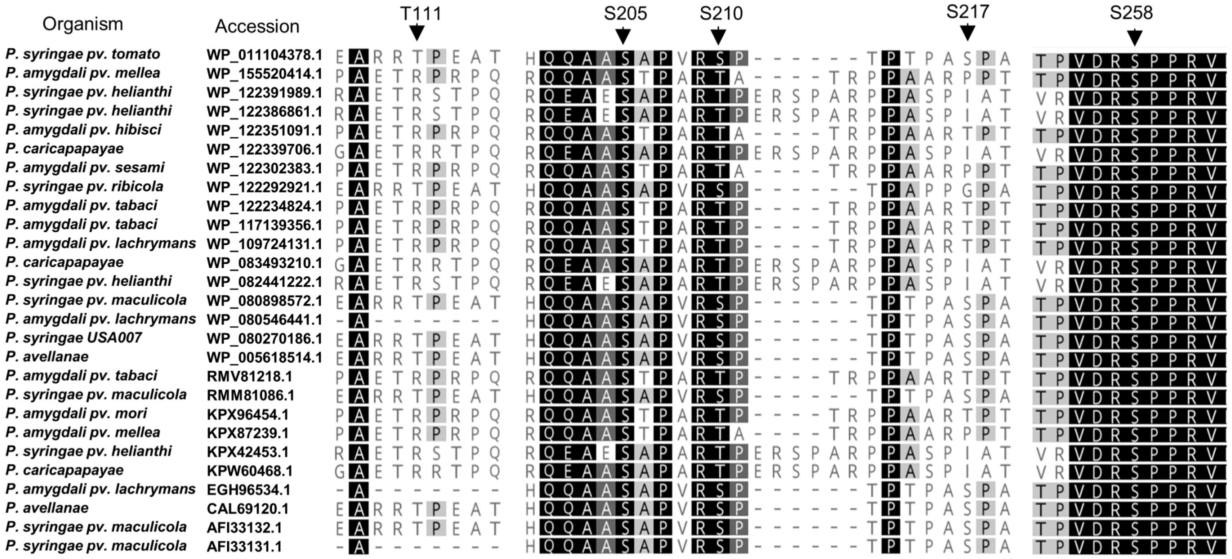


**Supplemental Figure 5. Conservation of phosphorylated residues in different AvrPtoB homologs. Related to Figure 2.**

AvrPtoB phosphorylated residues identified by mass spectrometry were analyzed for their conservation across 27 AvrPtoB homologs from *P. syringae*. The translated sequence of avrptoB were used in BLAST searches against the NCBI nr database on January 5th, 2020 (*Pseduomonas syringae* group, taxid: 136849; default parameters otherwise) to identify homologs. Hits were filtered based on a minimum 65% sequence identity, a minimum 85% query coverage and an E-value of 0. Sequences were downloaded and aligned in Genious using Genious alignment (default parameters).The percentage of identity is shown by gradient white to black (white means less than 60% similar, gray means 60-80% similar, dark gray means 80-100%, and black means 100% similar). AvrPtoB phosphorylation sites identified *in vivo* are shown with arrows.

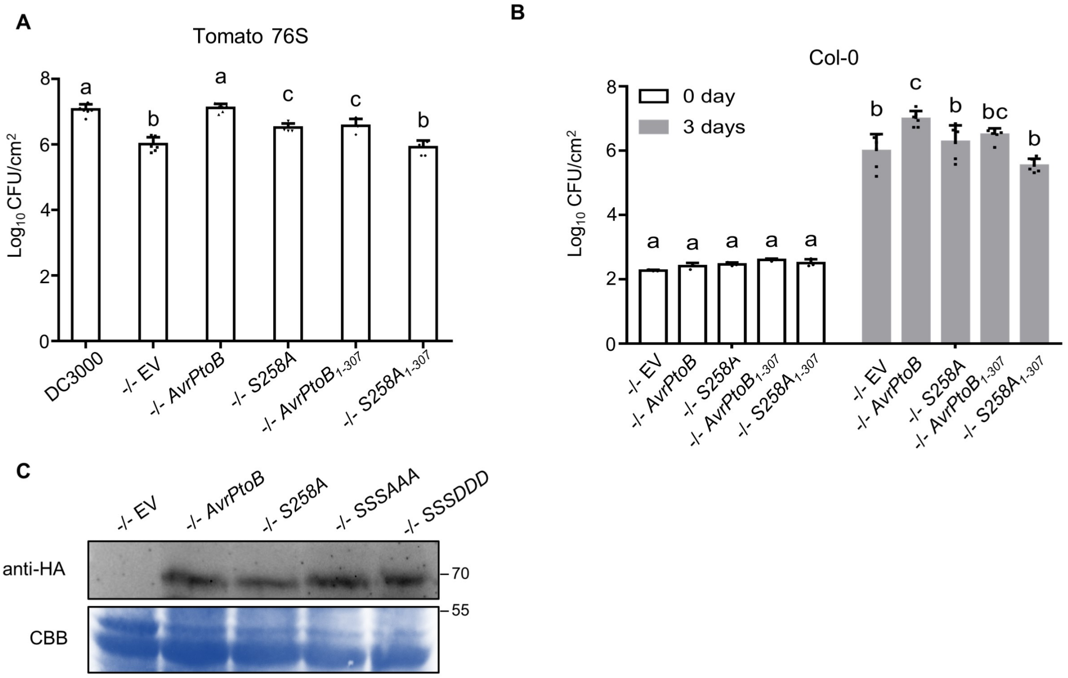


**Supplemental Figure 6. AvrPtoB S258 is required for virulence in tomato and *Arabidopsis*. Related to Figure 3.**

(A) Bacterial populations in the susceptible tomato genotype Rio Grande 76S five days post-inoculation with *P. syringae* pv. *tomato* DC3000Δ*avrPto*Δ*avrPtoB* (-/-) variants. DC3000 -/- carrying empty vector (EV) or plasmids expressing wild-type *AvrPtoB*, $AvrPtoB_{S258A}$, $AvrPtoB_{1\text{-}307}$, and $AvrPtoB_{1\text{-}307S258A}$ were dip inoculated on tomato at a concentration of $2 \times 10^8$ CFU $mL^{-1}$ with 0.005% Silwet. $Log_{10}$ CFU/$cm^2$ = $log_{10}$ colony-forming units per $cm^2$ of leaf tissue. Data are means ± SD (n ≥ 6 plants). Different letters indicate significant differences (one-way ANOVA, Tukey's test, $p < 0.05$).
(B) Bacterial populations in *Arabidopsis* Col-0 three days post-inoculation with DC3000 -/- variants. DC3000 -/- variants as described in (A) were syringe infiltrated in Col-0 at a concentration of $2 \times 10^5$ CFU $mL^{-1}$. Data are means ± SD (n = 4 plants for day 0, n = 6 plants for day 3). Different letters indicate significant differences (two-way ANOVA, Tukey's test, $p < 0.05$).
(C) Expression of AvrPtoB-HA and its derivatives from DC3000 -/- . Bacteria were grown in minimal media at 18 °C. Anti-HA western blotting was performed to detect the expression of AvrPtoB in bacterial pellets.

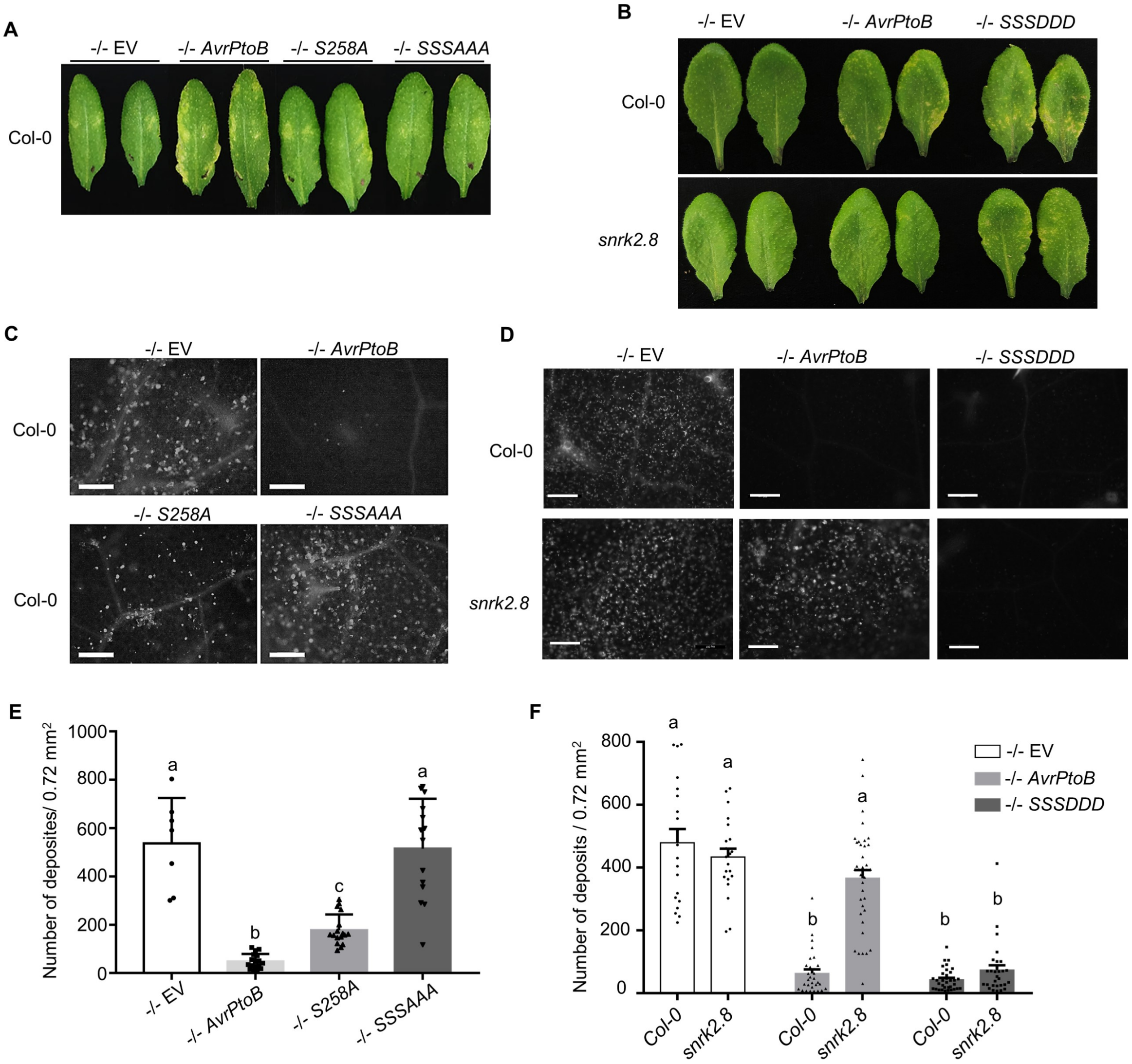


**Supplemental Figure 7. Phosphorylated AvrPtoB serine residues are required for virulence. Related to Figure 3.**

(A) Disease symptoms of *Arabidopsis* Col-0 after inoculation with *P. syringae* pv. *tomato* DC3000Δ*avrPto*Δ*avrPtoB* (-/-) variants. DC3000 -/- carrying empty vector (EV) or plasmids expressing wild-type *AvrPtoB*, $AvrPtoB_{S258A}$, or $AvrPtoB_{S205AS210AS258A}$ (*SSSAAA*) were syringe infiltrated into *Arabidopsis* Col-0 at a concentration of $2 \times 10^5$ CFU $mL^{-1}$. Disease symptoms were observed 4 days post-inoculation (dpi).
(B) Disease symptoms of *Arabidopsis* Col-0 after inoculation with *P. syringae* pv. *tomato* DC3000Δ*avrPto*Δ*avrPtoB* (-/-) variants. DC3000 -/- carrying empty vector (EV) or plasmids expressing wild-type *AvrPtoB*, or $AvrPtoB_{S205DS210DS258D}$ (*SSSDDD*). Plant were treated as described in (A).
(C) Callose deposition in *Arabidopsis* Col-0 after inoculation with -/- carrying EV, *AvrPtoB*, *S258A*, and *SSSAAA*. Leaves were inoculated with *P. syringae* at a concentration of $1 \times 10^8$ CFU $mL^{-1}$ and harvested 16h later. Leaves were stained by 1% aniline blue and imaged by fluorescence microscopy. Scale bar, 100 µm.
(D) Callose deposition in *Arabidopsis* Col-0 after inoculation with -/- carrying EV, *AvrPtoB*, and *SSSDDD*. Leaves were inoculated and callose visualized as described in (C).
(E) Quantification of callose deposits. Data are means ± SD (n = 8 images from 8 leaves for -/- EV, n = 18 images from 18 leaves for rest variants). Different letters indicate significant differences (one-way ANOVA, Tukey's test, $p < 0.05$).
(F) Quantification of callose deposits. Data are means ± SD (n = 21 images from 21 leaves for -/- EV, n = 30 images from 30 leaves for rest variants). Different letters indicate significant differences (one-way ANOVA, Tukey's test, $p < 0.05$).

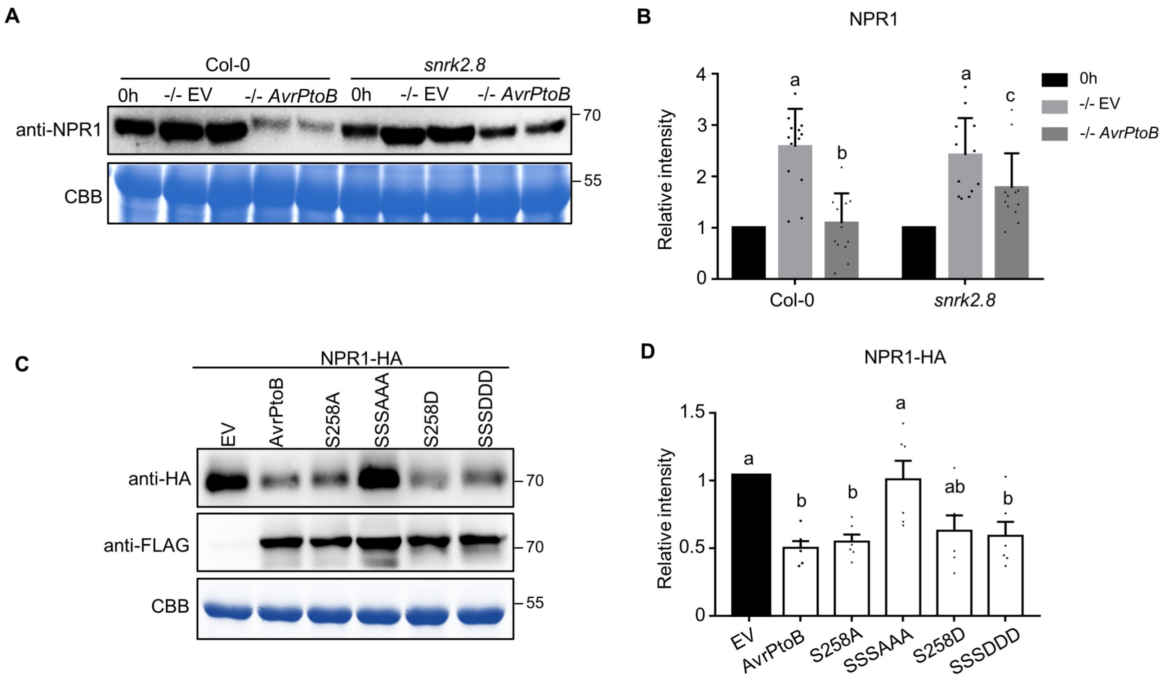


**Supplemental Figure 8. AvrPtoB mediated degradation of NPR1 requires the plant kinase SnrK2.8 and phosphorylated residues. Related to Figure 4.**

(A) NPR1 accumulation in *Arabidopsis* Col-0 and the *snrk2.8* knockout after inoculation with *P. syringae* pv. *tomato* DC3000Δ*avrPto*Δ*avrPtoB* (-/-) variants. DC3000 -/- carrying the empty vector (EV) or a plasmid expressing wild-type *AvrPtoB* were syringe infiltrated into Col-0 and *snrk2.8* at a concentration of $1 \times 10^8$ CFU $mL^{-1}$ and proteins extracted after four hours. Protein extracts were subjected to anti-NPR1 immunoblotting. Coomassie Brilliant Blue (CBB) staining shows equal protein loading.
(B) Quantification of NPR1-HA band intensity in (C). NPR1-HA bands intensities were quantified by Image Lab 6.0.1 (BIO-RAD). The values were normalized first by Rubisco bands and subsequently by the intensities of "EV" treatment. Data are means ± SD (n = 6 leaves). Different letters indicate significant differences (one-way ANOVA, Tukey's test, $p < 0.05$).
(C) NPR1 accumulation in the presence of AvrPtoB phosphorylation mutants in *N. benthamiana* after Agrobacterium-mediated transient expression. *35S::NPR1-HA* was co-expressed with FLAG-tagged Dex-inducible *AvrPtoB*, $AvrPtoB_{S258A}$, $AvrPtoB_{S205AS210AS258A}$ (*SSSAAA*), $AvrPtoB_{S258D}$ or $AvrPtoB_{S205DS210DS258D}$ (*SSSDDD*). The Dex inducible EV was used as a control. The expression of AvrPtoB-FLAG phosphorylation variants was induced by 15 µM DEX for 5 hours 24h post-Agrobacterium infiltration. Protein extracts were subjected to anti-HA and anti-FLAG immunoblotting. CBB staining shows protein loading.
(D) Quantification of NPR1 band intensity in (A). NPR1 bands intensities were quantified by Image Lab 6.0.1 (BIO-RAD). The values were normalized first by Rubisco bands and subsequently by the intensities of "0h" bands. Data are means ± SD (n = 14 leaves). Different letters indicate significant differences (one-way ANOVA, Tukey's test, $p < 0.05$).

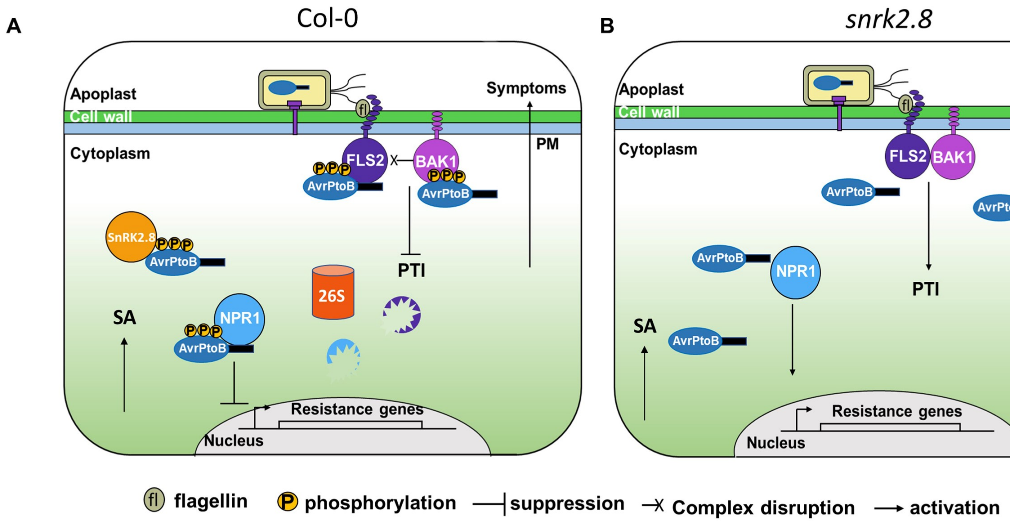


**Supplemental Figure 9. Model of AvrPtoB phosphorylation by SnRK2.8. Related to Figure 1- 4 and Supplemental figure 7, 8.**

(A) SnRK2.8 phosphorylates AvrPtoB in *Arabidopsis* Col-0 and the phosphorylated AvrPtoB mediates degradation of FLS2/NPR1 and inhibition of FLS2 complex formation.
(B) AvrPtoB virulence function is lost in *snrk2.8* knockout.

**Supplemental Table 1. The phosphorylation of bacterial type III effectors and their phosphorylation sites are identified by mass spectrometry. Related to Figure 1.**

Data were from the references in the table. Red indicates the phosphorylated residue.

| Effector | Pathogen | Phosphorylation site | Host kinase | Required for virulence | Required for avirulence | Reference |
|---|---|---|---|---|---|---|
| AvrPto | *P. syringae* | NPNG$S_{149}$IRMTL | unknown | Yes | Yes | Anderson et al., 2006<br>Yeam et al., 2009 |
| AvrPtoB | *P. syringae* | PVDR$S_{258}$PPRVNQR | unknown | Yes | Yes | Xiao et al., 2007 |
| HopQ1 | *P. syringae* | PVLERSK$S_{51}$APAL | SnRK (*in vitro*) | Yes | unknown | Li et al., 2013<br>Giska et al., 2013 |
| AvrB | *P. syringae* | unknown | RIPK (*in vitro*) | unknown | Yes | Desveaux et al., 2007<br>Chung et al., 2014 |
| AvrBsT | *X. campestris* | unknown | CaPIK1 (*in vitro*) | unknown | Yes | Kim et al., 2014 |
| XopN | *X. campestris* | APPRREHV$S_{688}$APSSP | unknown | Yes | unknown | Taylor et al., 2012 |
| XopE2 | *X. euvesicatoria* | RGRFRQP$T_{66}$LQPH<br>LSNGRSATY$S_{133}$SLSY<br>AQPI$S_{334}$PRTTAA | unknown | unknown | unknown | Dubrow et al., 2018 |
| NopL | *Rhizobium* | RSGPSQAGL$S_{139}$PSAT<br>SPSATPLLNP$S_{148}$PPPH<br>LTAERGR$S_{198}$PQPS | SIPK (*in vitro*) | Yes | unknown | Zhang et al., 2011a<br>Ge et al., 2016 |
| NopP | *Rhizobium* | unknown | unknown | Yes | unknown | Skorpil et al., 2005 |

### Supplemental Table 2. Related to Figure 2 and supplemental figure 4.

AvrPtoB was expressed in Col-0 and *snrk2.8* protoplasts and total proteins were subjected to anti-GFP IP followed by tryptic digestion. Phosphorylated peptides were detected by LC-MS/MS with the parallel reaction monitoring method. Peptide phosphorylation ratios were determined using Skyline software. Phoshorylation intensity is reported for residue T111 as no unphosphorylated peptides were detected by MS. Data are means ± SE of three biological replicates (separate transfections).

| | **Phosphorylation ratio** | | **SE** | |
|---|---|---|---|---|
| **Phosphorylated residue** | **Col-0** | ***snrk2.8*** | **Col-0** | ***snrk2.8*** |
| **S205** | 93.88 | 10.20 | ± 46.05 | ± 5.12 |
| **S210** | 173.85 | 17.90 | ± 85.40 | ± 8.81 |
| **S258** | 40263.92 | 4272.50 | ± 25060.68 | ± 2455.38 |
| **S217** | 0.01 | 0.01 | ± 0.001 | ± 0.003 |
| | **Phosphorylation intensity** | | **SE** | |
| | **Col-0** | ***snrk2.8*** | **Col-0** | ***snrk2.8*** |
| **T111** | 3880000 | 13500000 | ± 2837677.1 | ± 2478126.5 |

**Supplemental Table 3. Primers used in this study. Related to Methods.**

| Name | Sequence 5' to 3' | Purpose |
|---|---|---|
| SnRK1.1-LP | CACCATGTTCAAACGAGTAGATGAGTTT | Full-length SnRK1.1 CDS |
| SnRK1.1-RP | GAGGACTCGGAGCTGAGCAAG | |
| SnRK1.2-LP | CACCATGGATCATTCATCAAATAGATTTGGC | Full-length SnRK1.2 CDS |
| SnRK1.2-RP | GATCACACGAAGCTCTGTAAGAAAGG | |
| SnRK1.3-LP | CACCATGGATGGATCATCGGAAAAAAC | Full-length SnRK1.3 CDS |
| SnRK1.3-RP | GAGGACACCTAGCTCTCTGAGAAACG | |
| SnRK2.1-LP | CACCATGGACAAGTATGACGTTGTC | Full-length SnRK2.1 CDS |
| SnRK2.1-RP | AGCTTTGTCAGACTCTTGACAAGACTG | |
| SnRK2.3-LP | CACCATGGATCGAGCTCCGG | Full-length SnRK2.3 CDS |
| SnRK2.3-RP | GAGAGCGTAAACTATCTCTCCGCTAC | |
| SnRK2.4-LP | CACCATGGACAAGTACGAGCTGG | Full-length SnRK2.4 CDS |
| SnRK2.4-RP | ACTTATTCTCACTTCTCCACTTGCGTG | |
| SnRK2.5-LP | CACCATGGACAAGTATGAGGTTGTGAA | Full-length SnRK2.5 CDS |
| SnRK2.5-RP | AGCTTTGGGAGGCTCTTGACAAGAATG | |
| SnRK2.6-LP | CACCATGGATCGACCAGCAGTGA | Full-length SnRK2.6 CDS |
| SnRK2.6-RP | CATTGCGTACACAATCTCTCCGCTACTG | |
| SnRK2.7-LP | CACCATGGAGAGATACGACATCTTAAGAG | Full-length SnRK2.7 CDS |
| SnRK2.7-RP | TAGAGCACATACGAAATCACCATTTCTC | |
| SnRK2.8-LP | CACCATGGAGAGGTACGAAATAGTG | Full-length SnRK2.8 CDS |
| SnRK2.8-RP | CAAAGGGGAAAGGAGATCAGCG | |
| SnRK2.9-LP | CACCATGGAGAAGTATGAGATGGTG | Full-length SnRK2.9 CDS |
| SnRK2.9-RP | TGCGTAATCATCATACCATTCTTCATCA | |
| SnRK2.10-LP | CACCATGGACAAGTACGAGCTTGTTAA | Full-length SnRK2.10 CDS |
| SnRK2.10-RP | ACTGACTCGGACTTCTCCCATG | |
| SnRK3.1-LP | CACCATGGAGAAGAAAGGATCTGTGT | Full-length SnRK3.1 CDS |
| SnRK3.1-RP | GTGCCAAGCTAATACAAAGTCGATCAA | |
| SnRK3.2-LP | CACCATGGAGAACAAACCAAGTGTATT | Full-length SnRK3.2 CDS |
| SnRK3.2-RP | TGATGGTTCTTGCTCTCCTTGTTCATC | |
| SnRK3.6-LP | CACCATGGATAAAAACGGCATAGTTT | Full-length SnRK3.6 CDS |
| SnRK3.6-RP | ATGTATCACTTCAATCTTCTCATTGTTGC | |
| SnRK3.7-LP | CACCATGGCTCAAGTACTATCTACAC | Full-length SnRK3.7 CDS |
| SnRK3.7-RP | CTGTTCAATTTCAGGTGGCAAACA | |
| SnRK3.8-LP | CACCATGGAAAATAAGCCAAGTGTTTTG | Full-length SnRK3.8 CDSS |
| SnRK3.8-RP | AAACTTCAATGGTTCTTCCTGTTCTTG | |
| SnRK3.9-LP | CACCATGGCGGAGAAAATCACG | Full-length SnRK3.9 CDS |
| SnRK3.9-RP | TTCAGTGTCAGACGGCAAGAAAGAAATG | |

| Name | Sequence 5' to 3' | Purpose |
|---|---|---|
| SnRK3.10-LP | CACCATGGAATCACTTCCCCAG | Full-length SnRK3.10 CDS |
| SnRK3.10-RP | CATGATGTCATTGTGCCATGAAAGAAC | |
| SnRK3.11-LP | CACCATGACAAAGAAAATGAGAAGAGTGG | Full-length SnRK3.11 CDS |
| SnRK3.11-RP | AAACGTGATTGTTCTGAGAATCTCTGAC | |
| SnRK3.12-LP | CACCATGAGTGGAAGCAGAAGGAAG | Full-length SnRK3.12 CDS |
| SnRK3.12-RP | TTGCTTTTGTTCTTCAGCGGCTG | |
| SnRK3.13-LP | CACCATGGTGGTAAGGAAGGTG | Full-length SnRK3.13 CDS |
| SnRK3.13-RP | ACGTCTTTTACTCTTGGCCTTGGTG | |
| SnRK3.14-LP | CACCATGGTCGGAGCAAAACC | Full-length SnRK3.14 CDS |
| SnRK3.14-RP | AGCAGGTGTAGAGGTCCAGAAAATG | |
| SnRK3.15-LP | CACCATGGTAGATTCTGACCCGG | Full-length SnRK3.15 CDS |
| SnRK3.15-RP | CGACGTCGTATGTACTTGAGTTGGTTC | |
| SnRK3.16-LP | CACCATGGTGAGAAGGCAAGAG | Full-length SnRK3.16 CDS |
| SnRK3.16-RP | AGTTACTATCTCTTGCTCCGGCGAG | |
| SnRK3.17-LP | CACCATGTTGATCCCCAACAAAAAATTA | Full-length SnRK3.17 CDS |
| SnRK3.17-RP | CTTTGCTGTTTCTTTCTTAACTTCGTTAT | |
| SnRK3.19-LP | CACCATGTCTTTTACAATTCCTAGACTG | Full-length SnRK3.18 CDS |
| SnRK3.19-RP | CGGTTTGTCAGGAACTTTATAAACCG | |
| SnRK3.20-LP | CACCATGGCTCAAGCCTTGG | Full-length SnRK3.20 CDS |
| SnRK3.20-RP | TTCAGTATCAGATGGCAAATACAATGCTTC | |
| SnRK3.21-LP | CACCATGGTGATAAAGGGAATGCG | Full-length SnRK3.21 CDS |
| SnRK3.21-RP | CGCTAAAAGCTCCTGTACTTGTGATG | |
| SnRK3.22-LP | CACCATGCCAGAGATCGAGATTGC | Full-length SnRK3.22 CDS |
| SnRK3.22-RP | AATAGCCGCGTTTGTTGACGACG | |
| SnRK3.23-LP | CACCATGGCTTCTCGAACAACG | Full-length SnRK3.23 CDS |
| SnRK3.23-RP | TGTCGACTGTTTTGCAATTGTCCG | |
| SnRK3.24-LP | CACCATGGAGGAAGAACGG | Full-length SnRK3.24 CDS |
| SnRK3.24-RP | ACAATCCTCGGAAGAAGTGTTATTATTA | |
| SnRK3.25-LP | CACCATGGGATCCAAACTTAAACTTTAC | Full-length SnRK3.25 CDS |
| SnRK3.25-RP | GCAGTCACTACCAGAATTTTCATCAC | |
| CPK2ΔC-LP | CACCATGGGTAATGCTTGCGTT | CPK2 C-terminal deleted CDS |
| CPK2ΔC-RP | CACACCGTCAATCTGTACCCAT | |
| CPK3ΔC-LP | CACCATGGGCCACAGACACAGC | CPK3 C-terminal deleted CDS |
| CPK3ΔC-RP | CTCCCCATCTTCTCTAATCCACGG | |
| CPK4ΔC-LP | CACCATGGAGAAACCAAACCCT | CPK4 C-terminal deleted CDS |
| CPK4ΔC-RP | AGCATGTTCATCAACAATCCAAG | |
| CPK5ΔC-LP | CACCATGGGCAATTCTTGCC | CPK5 C-terminal deleted CDS |
| CPK5ΔC-RP | AACACCATTCTCACAGATCCATG | |

| Name | Sequence 5' to 3' | Purpose |
|---|---|---|
| CPK6ΔC-LP | CACCATGGGCAATTCATGTCGT | CPK6 C-terminal deleted CDS |
| CPK6ΔC-RP | AACTCCATTCTCACAGATCCATG | |
| CPK10ΔC-LP | CACCATGGGTAACTGTAACGCC | CPK10 C-terminal deleted CDS |
| CPK10ΔC-RP | TTTCTTTGCATTCTGTATCCATGGG | |
| CPK11ΔC-LP | CACCATGGAGACGAAGCCAAAC | CPK11 C-terminal deleted CDS |
| CPK11ΔC-RP | TGCTTGTTCATCGACAATCCATG | |
| CPK13ΔC-LP | CACCATGGGAAACTGTTGCAGA | CPK13 C-terminal deleted CDS |
| CPK13ΔC-RP | TTTCTTTGCGTTCTGAATCCATGG | |
| CPK26ΔC-LP | CACCATGAAGCACAGCGGTGG | CPK26 C-terminal deleted CDS |
| CPK26ΔC-RP | AACTCCATTTTCACAGATCCAAGG | |
| SnRK1.1 KA-LP | CATAAGGTTGCTATCGCGATCCTCAATCGTC | Generate SnRK1.1 kinase dead mutant |
| SnRK1.1 KA-RP | GACGATTGAGGATCGCGATAGCAACCTTATG | |
| SnRK2.6 KA-LP | CTTGTTGCTGTTGCATATATCGAGAG | Generate SnRK2.6 kinase dead mutant |
| SnRK2.6 KA-RP | CTCTCGATATATGCAACAGCAACAAG | |
| SnRK2.8 KA-LP | GAGCTTTTCGCTGTTGCGTTCATCGAGCGAG | Generate SnRK2.8 kinase dead mutant |
| SnRK2.8 KA-RP | CTCGCTCGATGAACGCAACAGCGAAAAGCTC | |
| CPK4ΔC KA-LP | CTAATTACGCTTGCGCATCAATCCCAAAAC | Generate CPK4ΔC kinase dead mutant |
| CPK4ΔC KA-RP | GTTTTGGGATTGATGCGCAAGCGTAATTAG | |
| CPK5ΔC KA-LP | GACTACGCTTGTGCGTCAATATCCAAG | Generate CPK5ΔC kinase dead mutant |
| CPK5ΔC KA-RP | CTTGGATATTGACGCACAAGCGTAGTC | |
| AvrPtoB-LP | CACCATGGCGGGTATCAATGGAGC | Clone Full-length AvrPtoB |
| AvrPtoB-RP | GGGGACTATTCTAAAAGC | |
| AvrPtoB 307-RP | TACATGTCTTTCAAGGGCCGTG | Clone $AvrPtoB_{1-307}$ |
| AvrPtoB S258A -LP | CCGGTCGACAGGGCCCCGCCACGCG | Generate AvrPtoB S258A mutant |
| AvrPtoB S258A-RP | CGCGTGGCGGGGCCCTGTCGACCGG | |
| AvrPtoB S258D-LP | CCGGTCGACAGGGACCCGCCACGCG | Generate AvrPtoB S258D mutant |
| AvrPtoB S258D-RP | CGCGTGGCGGGTCCCTGTCGACCGG | |
| AvrPtoB S205S210A-LP | CAACAGGCGGCGGCAGCGCCAGTGAGGGCGCCCACGCCAAC | Generate AvrPtoB S205AS210AS258A mutant |
| AvrPtoB S205S210A-RP | GTTGGCGTGGGCGCCCTCACTGGCGCTGCCGCCGCCTGTTG | |
| AvrPtoB S205DS210D-LP | CAACAGGCGGCGGATGCGCCAGTGAGGGATCCCACGCCAAC | Generate AvrPtoB S205DS210DS258D mutant |
| AvrPtoB S205DS210D-RP | GTTGGCGTGGGATCCCTCACTGGCGCATCCGCCGCCTGTTG | |
| snrk2.8 SALK_073395-LP | ATTTTCCAAAGAGCTTTTCGC | snrk2.8 T-DNA line genotyping |
| snrk2.8 SALK_073396-RP | GGTGATAGGTTTCCGAGCTTC | |

**Supplemental Table 4. Isolation list for PRM. Related to Figure 2.**
Modified and unmodified peptide sequences used to quantify phosphorylation by PRM are noted. CID = collision induced dissociation, m/z = mass to charge, z = charge state.

| Peptide sequence | Peptide modified sequence | m/z | z |
|---|---|---|---|
| AEARRTPEATADASAPR | AEARRT[+80]PEATADASAPR | 617.2899 | 3 |
| RAVHQQAASAPVR | RAVHQQAASAPVR | 464.2603 | 3 |
| RAVHQQAASAPVRSPTPTPASPAASSSGSSQR | RAVHQQAASAPVRSPTPTPASPAASSSGSSQR | 786.9028 | 4 |
| | RAVHQQAASAPVRSPTPTPASPAASSSGSSQR | 629.7237 | 5 |
| | RAVHQQAAS[+80]APVRSPTPTPASPAASSSGSSQR | 806.8944 | 4 |
| | RAVHQQAASAPVRS[+80]PTPTPASPAASSSGSSQR | 806.8944 | 4 |
| AVHQQAASAPVR | AVHQQAASAPVR | 617.8362 | 2 |
| | AVHQQAASAPVR | 412.2265 | 3 |
| AVHQQAASAPVRSPTPTPASPAASSSGSSQR | AVHQQAASAPVRSPTPTPASPAASSSGSSQR | 996.8343 | 3 |
| | AVHQQAASAPVRSPTPTPASPAASSSGSSQR | 747.8775 | 4 |
| | AVHQQAAS[+80]APVRSPTPTPASPAASSSGSSQR | 767.8691 | 4 |
| | AVHQQAASAPVRS[+80]PTPTPASPAASSSGSSQR | 1023.49 | 3 |
| | AVHQQAASAPVRS[+80]PTPTPASPAASSSGSSQR | 767.8691 | 4 |
| SPTPTPASPAASSSGSSQR | SPTPTPASPAASSSGSSQR | 886.9241 | 2 |
| | SPTPTPASPAASSSGSSQR | 591.6185 | 3 |
| | SPTPTPAS[+80]PAASSSGSSQR | 926.9073 | 2 |
| SSNTAASQTPVDRSPPR | SSNTAASQTPVDRSPPR | 590.9625 | 3 |
| | SSNTAASQTPVDRS[+80]PPR | 925.9233 | 2 |
| | SSNTAASQTPVDRS[+80]PPR | 617.6179 | 3 |
| | PSNTPPSNAPAPPPTGR | 828.411 | 2 |
| | PSNT[+80]PPSNAPAPPPTGR | 868.3942 | 2 |
| AALDPIASQFSQLR | AALDPIASQFSQLR | 757.9023 | 2 |
| | AALDPIASQFS[+80]QLR | 797.8855 | 2 |
